# Geometry of spiking patterns in early visual cortex: a Topological Data Analytic approach

**DOI:** 10.1101/2022.03.16.484568

**Authors:** Andrea Guidolin, Mathieu Desroches, Jonathan D. Victor, Keith P. Purpura, Serafim Rodrigues

## Abstract

In the brain, spiking patterns live in a high-dimensional space of neurons and time. Thus, determining the intrinsic structure of this space presents a theoretical and experimental challenge. To address this challenge, we introduce a new framework for applying topological data analysis (TDA) to spike train data and use it to determine the geometry of spiking patterns in the visual cortex. Key to our approach is a parameterized family of distances based on the timing of spikes that quantifies the dissimilarity between neuronal responses. We applied TDA to visually driven single-unit and multiple single-unit spiking activity in macaque V1 and V2. TDA across timescales reveals a common geometry for spiking patterns in V1 and V2 which, among simple models, is most similar to that of a low-dimensional space endowed with Euclidean or hyperbolic geometry with modest curvature. Remarkably, the inferred geometry depends on timescale, and is clearest for the timescales that are important for encoding contrast, orientation, and spatial correlations.

## Introduction

The broad goal of understanding the nature and function of neural activity entails the intermediate step of characterizing the space in which this activity lives. This is a challenging problem, even for small populations of neurons, because of the dimensional explosion and data limitations: it is simply not feasible to obtain sufficient experimental data to exhaustively sample the space of all possible spiking patterns [46]. For this reason, indirect methods are required. A promising strategy, applicable in a wide variety of contexts, is the persistent homology approach to topological data analysis (TDA) [22, 36, 37, 50, 10]. With TDA, an abstract notion of distance is used to compare observations, such as samples of neural activity. Based on these pairwise distances, a sequence of network graphs can be constructed, with successive graphs built by connecting the nodes at increasingly greater distances. These graphs are then associated with higher-dimensional topological objects called simplicial complexes, which are characterized via their Betti numbers, describing (intuitively speaking) the number of disconnected components, the number of holes, the number of “bubbles”, etc. The way that the Betti numbers change as a function of the distance criterion between the samples then provides a topological characterization of the space.

Here, we use this approach to analyze the activity of clusters of cortical neurons, with a specific focus on patterns of spiking activity in time. This requires confronting another issue, but one that the persistent homology approach is also well-suited to handle: even for a single neuron, activity is not a scalar quantity, but rather, a sample of a point process. Thus, it is possible to use the TDA machinery in a novel way, by applying it to a measure of distances between neural responses that is sensitive to temporal patterns within and across neurons: the family of spike distances introduced by Victor and Purpura [44]. While these distances do not correspond to Euclidean distances [4], they do satisfy the triangle inequality, which suffices for the TDA machinery.

While our approach shares the overall goal of a geometric characterization of neural activity with many other studies, there are some important differences in the questions we ask. Below we highlight the two main distinctions.

One central question is, what is the dimensionality of the response space? In previous studies [12, 11] dimensionality is defined as the dimension of the space that contains response trajectories, where a response trajectory is the evolution of average firing rates over time. Thus, these previous approaches would not distinguish between spiking activity that was well-described by a Poisson process, and spiking activity whose patterning required additional dimensions to describe. Our construction of the response space is quite different: it is sensitive to the patterns of spikes on individual trials. We find that there are substantial differences between the geometry of the recorded neural activity and their Poisson-like equivalents. Moreover, we find that the influence of spike patterning on geometry is maximal at the timescales that are important for neural coding of visual attributes such as contrast, orientation and spatial patterns [43].

Many recent studies have shown that the geometry of the responses “untangles” the key parameters of the stimulus, an important and novel finding for complex, naturalistic domains [31, 23, 26]. This is not a focus of the current study, as an untangling is to be anticipated for the stimulus set studied here: a family of artificial visual textures parameterized by local image statistics [42]. V1 and V2 neurons are selectively tuned to these statistics (see below), so decoding of the population will necessarily identify distinct directions (i.e., distinct population vectors) that capture the stimulus parameters.

Rather, our focus is on whether the intrinsic geometry of neural activity patterns depends on the stimulus parameters being encoded. To this end, our visual stimuli contained a range of types of statistical structures, including textures that varied in first-, second-, third-, and fourth-order local spatial correlations (see Materials and Methods) known to be present in natural images [27] and to be salient for human observers [45]. First- and second-order statistics control luminance and orientation content, respectively, and thus drive neurons in both V1 and V2, while third- and fourth-order statistics control aspects of local form that are primarily detected by V2 neurons [49]. Here, we separately analyze responses to many examples of textures that explore each of these statistics. Thus, we can determine whether the geometry of neural activity patterns depends on the kind of visual structure that is encoded, or perhaps on the cortical area (V1 vs. V2), or alternatively, has a more universal behavior – the latter suggesting that the spike patterning dynamics are endogenously-driven.

Applying TDA to spike distances carries with it a theoretical departure from most previous applications of the persistent homology approach to neural data. Provided that mean responses depend continuously on stimulus parameters, Euclidean distances between mean firing rates or distances derived from correlations [22, 13, 14], which are effectively dot products in a vector space, will always lead to a manifold (though not necessarily one of low dimension). The reason is that these distance will map neural activity to a smooth and connected subset of a vector space. Spike metrics have a discrete component and consequently are fundamentally not Riemannian, so response geometry will typically not constitute a manifold – but, as we show, manifold structure is not needed to interpret the results of TDA.

Finally, we find that a novel way of passing from distance measures to graphs – linking nodes according to greatest dissimilarity rather than greatest similarity (using a “decreasing filtration” rather than the standard “increasing filtration”) – sharpens the distinction between the experimental data and candidate geometrical models.

## Results

The starting point of our analysis is 28 datasets of single-unit spiking activity recorded from areas V1 and V2 of the visual cortex of anesthetized macaques during stimulation with visual patterns drawn from a high-dimensional space of visual textures [42]. Each dataset is a recording at a separate recording site or in a different macaque, and at each site, the stimulus space is sampled along its 10 axes, corresponding to 10 independent and visually salient kinds of spatial correlation (Fig. 1). We sample each axis at four points: two levels of positive correlation and two levels of negative correlation (see Materials and Methods for details). 64 perceptually similar examples of each of these 40 texture types (10 axes, 4 sample points per axis) were presented in random order, with each stimulus lasting 320 ms. This sequence was repeated four times at each recording site. We refer to the neuronal responses to the 64 perceptually similar stimuli obtained in this way in one repeat as a “collection”; thus, there were 160 collections (40 texture types, 4 repeats) in each dataset. For our analysis, we selected the 80 collections with the smallest number of empty responses from each dataset. Depending on the dataset, the responses consisted of the spike trains of one or more isolated neurons. If more than four neurons were simultaneously recorded, we considered only the four neurons with the highest firing rates. We then applied TDA to characterize the topology of the firing patterns in each of the collections, and summarize the resulting characterizations to identify their consistent features.

**Figure 1:**
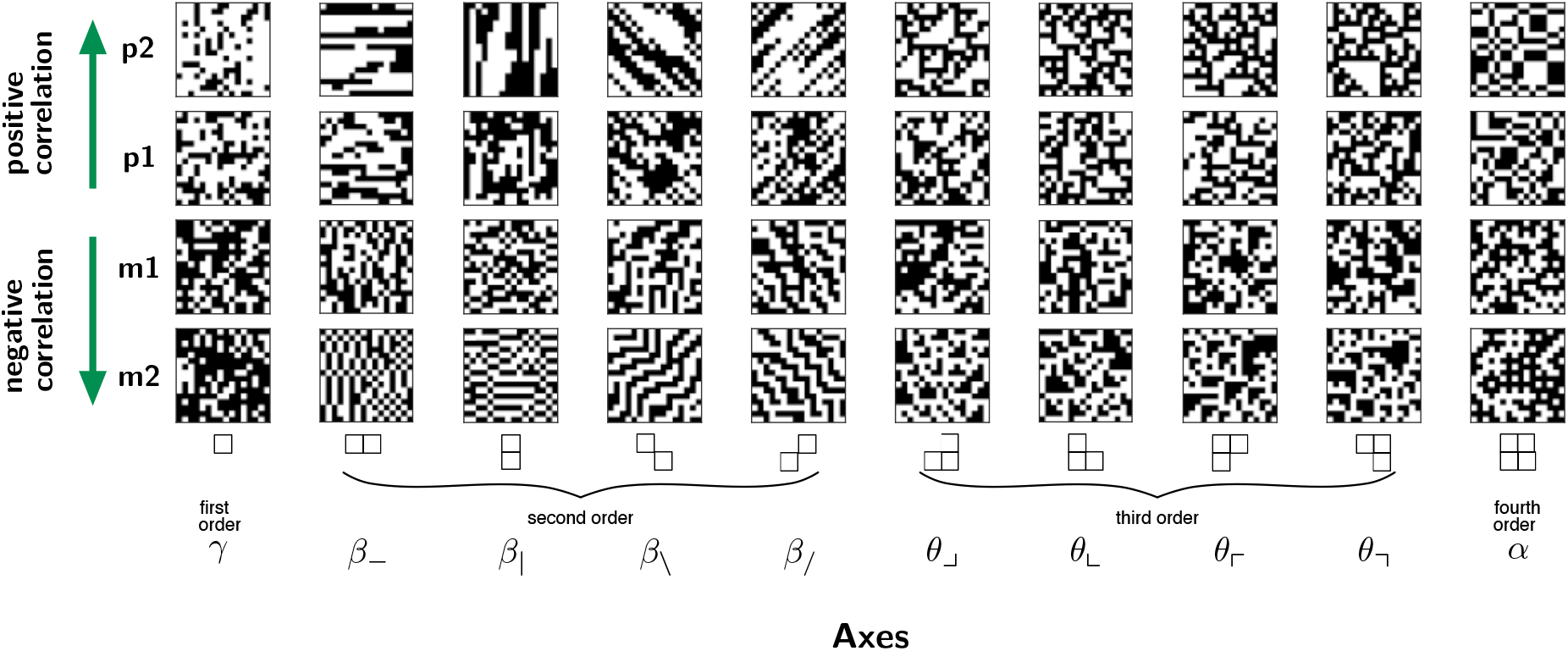
The space of visual stimuli. The figure shows examples of stimuli, randomly drawn from the 10 coordinate axes (*γ*, …, *α*) of a mathematically-defined stimulus space, with 4 correlations strengths (m2, m1, p1, p2) along each axis. Each stimulus is a 16×16 black and white checkerboard-like pattern. The coordinate axes of the stimulus space specify the type of the multipoint correlation within 2 2 clusters of checks, and are divided into four classes according to the order of correlation [45]: first-order (*γ*), second-order (*β*_−_, *β*_|_, *β*_*/*_, *β*_*/*_), third-order (*θ*_⌟_, *θ*_⌞_, *θ*_⌜_, *θ*_⌝_), and fourth-order (*α*). The correlation strengths, which are the coordinates along these axes, can have positive or negative sign. They determine the frequency with which local configurations of 1, 2, 3, or 4 checks (diagrammed at the bottom of each column of textures) appear in the pattern with even vs. odd parity. Specifically, a correlation strength of *c* means that instances of configurations with an even number of black checks occur with probability (1 + *c*)/2, and configurations with an odd number of black checks occur with probability (1−*c*) 2. To generate the visual stimuli used in the experiment, each axis of the space is sampled at four correlation strengths: high negative (m2), low negative (m1), low positive (p1) and high positive (p2). Correlation values were ±0.2 and ±0.4 for first-order correlations, and ±0.4 and ±0.8 for second-, third-, and fourth-order correlations.

The TDA method we used follows very closely the framework proposed by [22], but we started from a different way of endowing a set of neuronal responses with a notion of distance. Specifically, we used the multineuron Victor-Purpura distance [43, 3] to define the distances between all pairs of neuronal responses. This choice to employ a distance between spike trains rather than the correlations (based for example on firing rate) between neuronal responses to quantify (dis)similarity is motivated by several considerations. The main consideration is that we wanted to focus on spiking patterns in neuronal data that is relatively sparse (typically only a few spikes in the 320 ms response period). Information about spiking patterns would be lost in averages over multiple stimulus presentations, or after a smoothing procedure to estimate time-varying firing rates from single trials. Secondarily, use of the Victor-Purpura distance allows us to build on previous studies of the visual cortex that employ other data analytical techniques. We note that the generality of the TDA approach extends seamlessly to metrics such as the Victor-Purpura distance, as it does not assume a vector space embedding.

The Victor-Purpura distance between spike trains (see Materials and Methods) is defined as the minimum cost of transforming one spike train into the other via addition or deletion of spikes, shift of spike times, or change in the neuron of origin of the spikes. Each editing move is associated with a cost, which depends on a parameter *q*, controlling the relevant timescale for shifts of spike times, or on a parameter *k*, controlling the sensitivity to the neuron of origin of each spike. The family of distances defined in this way ranges from measures of dissimilarity that ignore spike timing altogether (*q* = 0), measuring distance based only on spike count, to measures that consider spike timing meaningful at an arbitrarily high level of precision (*q* → ∞). Similarly, *k* = 0 corresponds to ignoring the neuron of origin, while *k* = 1 corresponds to considering a change in the neuron of origin equivalent to the insertion of a spike. We carried out TDA for a grid of values of these two parameters: *q=*1, 2, 5, 10, 20, 50, 100, 200 (sec^−1^) and *k* = 0, 1. The resulting grid covered the range found relevant to neural coding in previous studies [43, 3].

For each choice of parameters of the Victor-Purpura distance we computed topological summaries via Betti curves [22] (Fig. 2) and characterized relevant features of the topological space by computing mean Betti curves averaged over the 80 collections drawn from of each dataset. The procedure for construction of the Betti curves is detailed in Materials and Methods and summarized here. We first create a sequence of graphs from each collection; in each graph, the nodes are the individual responses in the collection, and the edges are determined by the dissimilarities (Fig. 2A). As in the standard implementation of TDA, each graph is built by connecting nodes whose dissimilarities are less than a given ceiling. Thus, the graph sequence begins with a ceiling of 0 (all nodes disconnected) and terminates at a ceiling equal to the maximum distance (all nodes connected), a graph sequence known as the *increasing filtration*, and used in the *clique topology* method of [22]. Here, we also create a graph sequence by adding in the edges in decreasing order (the *decreasing filtration*), like in [32]. In each filtration, as the edges of the graph fill in, its topology can be characterized by the number of enclosed “holes” of different dimensions: the number of 1-dimensional tunnels, 2-dimensional voids and 3-dimensional “cavities”, known as Betti numbers and denoted by *β*_1_, *β*_2_ and *β*_3_ respectively. These are computed at each step of the filtration, and thus, the Betti numbers are functions of the edge density *ρ*, the fraction of potential edges that have been filled in: *β*_1_ (*ρ), β*_2_ (*ρ)* and *β*_3_ (*ρ)*. These quantities, which we computed over the range from *ρ* = 0 to 0.6 and then averaged over the 80 collections in a dataset, are shown in Fig. 2B.

**Figure 2:**
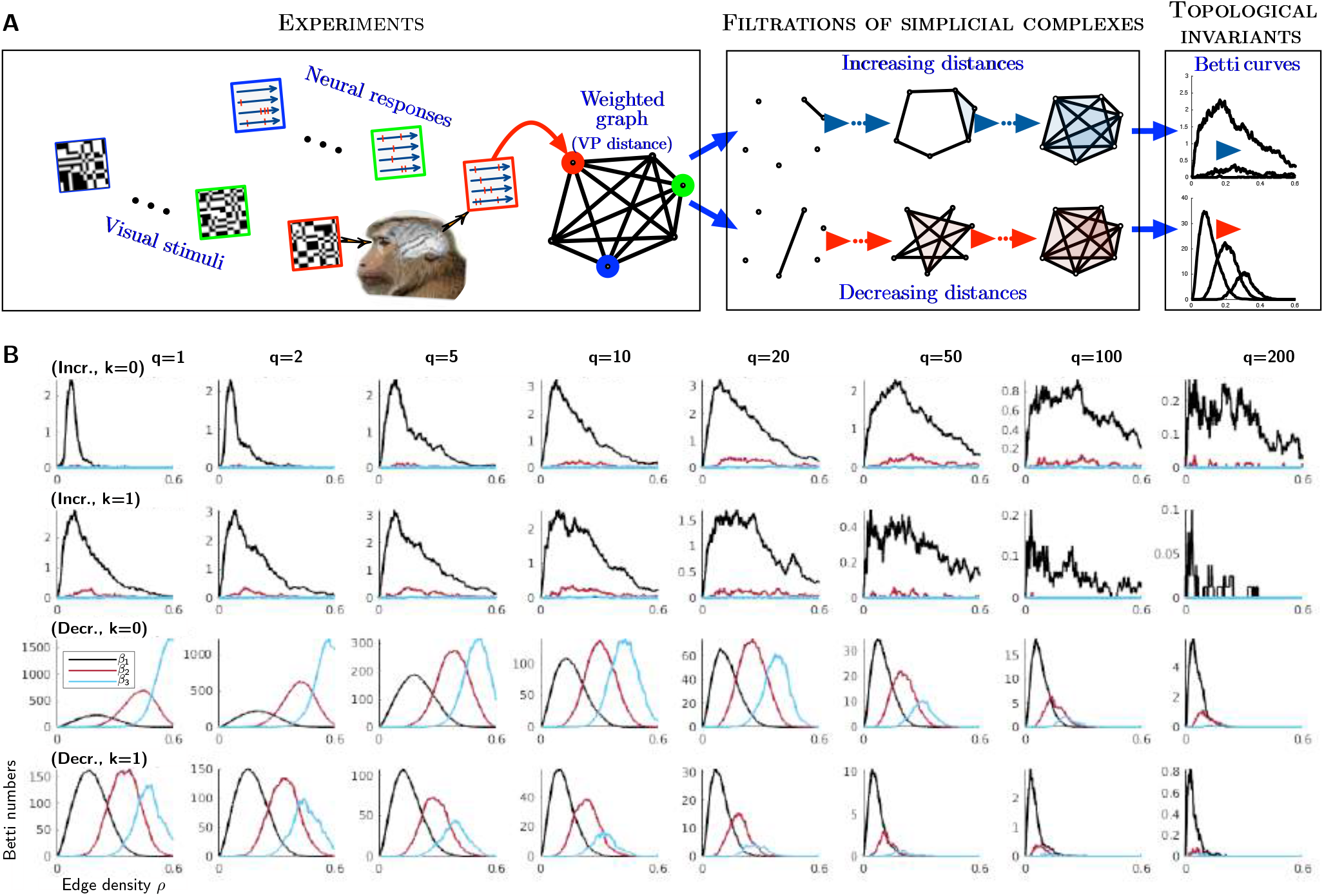
A. Pipeline for extracting topological summaries from the experimental data. Left: For each data set, neuronal responses (single-unit or up to four simultaneously-recorded units) are grouped into collections of 64 responses of duration 320 ms each. Each collection consists of responses to 64 different examples of textures defined by one of 10 texture parameters. The responses in each collection become points (colored balls) in a state space, whose distances, summarized by the weighted graph, are determined by the Victor-Purpura metric. Center: The weighted graph is associated with a filtration of simplicial complexes, using the clique topology method, by successively adding edges of the graph according to their weights, either increasingly or decreasingly. Right: Betti curves are computed from each filtration. Betti curves are a topological descriptor that captures how the number of topological voids of different dimensions (holes, “bubbles”, etc.) depends on the proportion of added edges in the filtration process. **B. Average Betti curves of one dataset**. The average Betti curves for *β*_1_-*β*_3_ over the 80 collections of one dataset (L7301TT6), containing recordings of spiking activity of 4 neurons in layer 5 of area V2, are displayed for increasing and decreasing filtrations, and for each value of the parameters *q* (timescale, sec^−1^) and *k* (sensitivity to neuron of origin) of the Victor-Purpura distance. The two top (respectively, bottom) rows correspond to the increasing (resp., decreasing) filtration method. The odd (resp., even) rows correspond to *k* = 0 (resp., *k* = 1). In the increasing filtration plots, the Betti curves for *β*_3_ are close to zero.

Note that while the Betti curves are sensitive to the geometry and topology of the metric space of data points [22, 37, 50] they are invariant under monotonic transformations of the distance [22]. This means that they are sensitive only to the relative ordering of all distances.

Below, we will focus on the mean Betti curves from the 80 collections with the highest firing rates at each recording site, but first we investigated whether the Betti curves had a substantial dependence on the two critical experimental variables that distinguished the collections: whether the recordings were in V1 vs. V2, and whether the spatial structure of the stimulus was low-order (first- and second-order) vs. high-order (third-or fourth-order) (see Fig. 1, Materials and Methods, and [45] for details and definitions). Low-order correlation structure is extracted by linear receptive fields and is already present in V1; third- and fourth-order correlation structure is primarily extracted in V2 [49].

To compare the topological characterizations of the collections of responses, we summarized the information in the Betti curves by two main features: the integral of the curves [22], referred to as integrated Betti values, and the position of their dominant peak on the *ρ*-axis, quantified by their center of mass (see Materials and Methods for details). Fig. 3 shows that the integrated Betti values of the collections recorded in areas V1 and V2 of the visual cortex have similar distributions. A remarkable consistency can be observed across all values of the parameters of the Victor-Purpura distance, and for both the increasing and decreasing filtration. The distributions of the center of mass of the Betti curves (Fig. S2) also show no dependence on recording area.

**Figure 3:**
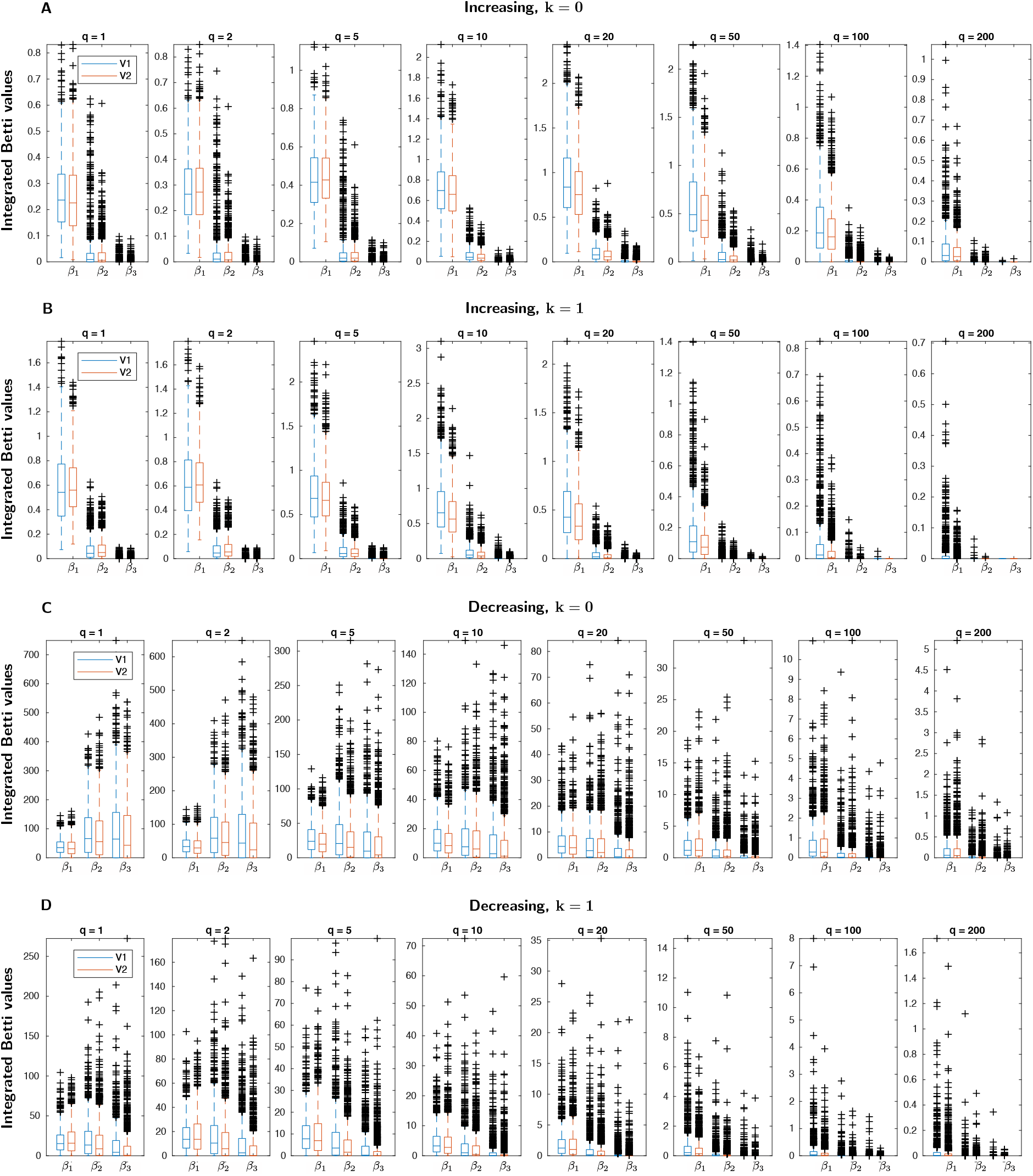
Comparison of integrated Betti values for data recorded in V1 vs. V2. Integrated Betti values (i.e., the values of the integrals of the Betti curves) for *β*_1_-*β*_3_ of the experimental data are shown divided into two groups, according to whether the data were recorded in area V1 (blue) or V2 (red) of the visual cortex. To generate the distributions of integrated Betti values displayed in the figure, individual (non-averaged over the dataset) Betti curves of each collection of responses are considered. The four panels are for increasing (**A**,**B**) and decreasing (**C**,**D**) filtrations, and for *k* = 0 (**A**,**C**) and *k* = 1 (**B**,**D**). Each panel shows the distribution of integrated Betti values for all values of the timescale parameter *q* of the Victor-Purpura distance.

Furthermore, the distributions of integrated Betti values (Fig. 4) and center of mass (Fig. S3) do not depend on whether the visually-salient structure of the textures is driven by first- and second-order spatial correlations, which lead to differences in luminance and spatial frequency that is already manifest in the cortical input, or by third- and fourth-order spatial correlations, which require intracortical calculations to extract [49].

**Figure 4:**
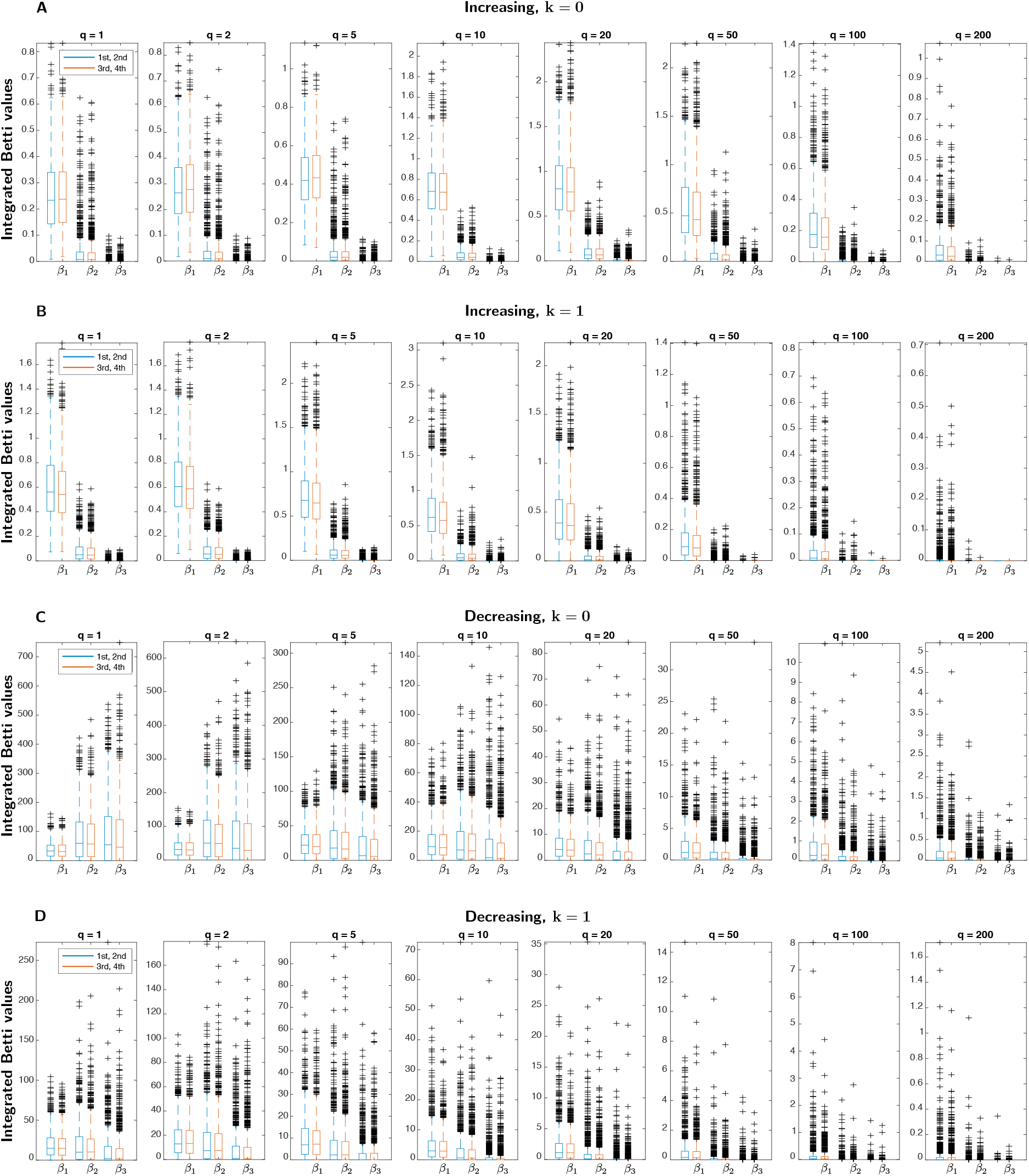
Comparison of integrated Betti values for responses elicited by low- and high-order spatial correlations. Integrated Betti values for *β*_1_-*β*_3_ of the experimental data are shown divided into two groups, according to whether neuronal responses are driven by visual stimuli with first- and second-order correlations (axes *γ, β* _−_, *β*_|_, *β ∖, β* of the stimulus space described in [45] and summarized in Fig. 1) or third- and fourth-order correlations (axes *θ* _*⌟*_, *θ* _*⌞*_, *θ* _*⌜*_, *θ* _*⌝*_and *α*). The distributions of integrated Betti values, determined as in Fig. 3, are respectively shown in blue and red. High-order (third- and fourth-order) correlations are extracted primarily in V2 [49]; low-order (first- and second-order) correlations can be extracted by the spatial filtering of retinal processing. To generate the distribution of integrated Betti values, individual (non-averaged over the dataset) Betti curves of each collection of responses are considered. The four panels are for increasing (**A**,**B**) and decreasing (**C**,**D**) filtrations, and for *k* = 0 (**A**,**C**) and *k* = 1 (**B**,**D**). Each panel shows the distribution of integrated Betti values for all values of the timescale parameter *q* of the Victor-Purpura distance.

Note that this analysis does not seek to determine whether the responses to one kind of texture differ from responses to another (whether they lie in different parts of a combined response space, e.g., by virtue of different firing rate profiles), but rather, to analyze the intrinsic geometry of each collection, via methods that focus on firing pattern. Our results up to this point suggest that this geometry is largely independent of the recording area, or the stimuli that drive the responses. Given the similarity of the results of TDA across recording areas and texture types, we pool these results in the further analysis below and now focus on the mean Betti curves over the 80 collections within each of the datasets.

To assess which aspects of the spike train data might be responsible for the shape of these mean Betti curves (e.g., Fig. 2B), we synthesized surrogate spike train data of four different types by perturbing specific aspects of the original experimental data. The description of the surrogate data is detailed in Materials and Methods and summarized here.

1. *Uniform resampling of spike times* (U): the spike times of each response are resampled uniformly in the response interval (0−320ms).
2. *Exchange resampling of spike times between collections* (EB): the spike times of each response are resampled uniformly, without replacement, in the set of spike times of the 80 collections of the dataset.
3. *Exchange resampling of spike times within collections* (EW): as in 2, but with the resampling restricted to the set of spike times of the collection to which the response belongs.
4. *Poisson generated spike trains* (P): each response is replaced with a sample of a Poisson process with the same firing rate.

In all cases, the spike times are resampled independently for each neuron that participates in a response. For each experimental dataset, we generated 20 surrogate spike datasets for each of the above four types, and computed Betti curves for these surrogates just as the curves were computed for the experimental data.

An example of this analysis is shown in Fig. 5. As is typical across the datasets, the Betti curves associated with the experimental data were often quite distinct from those of the surrogate sets. These differences were most prominent in the mid-range values of *q* (5 to 50 sec^−1^), and also more prominent for the decreasing filtration than for the increasing filtration. In addition, the Poisson surrogates usually show a greater divergence from the real data than the other surrogates. One possible reason for this is that, unlike the other surrogates, the Poisson surrogates do not preserve the number of spikes in each response.

**Figure 5:**
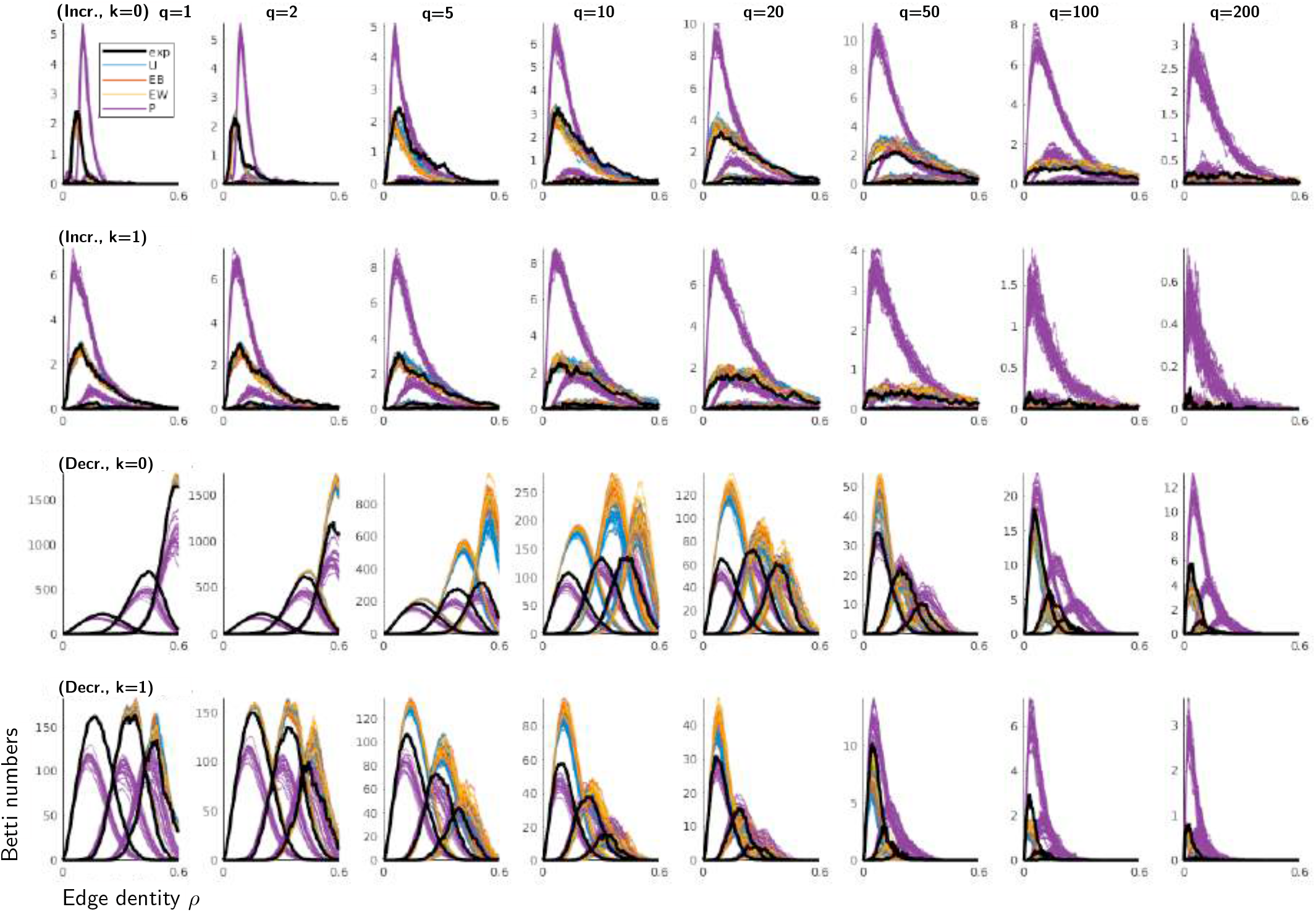
Average Betti curves of one dataset, and comparison with surrogate data. The figure superimposes on Fig. 2B the Betti curves of the 20 computations of the four types of surrogate data generated from the single dataset “L7301TT6”. The average Betti curves for *β*_1_-*β*_3_ of the experimental data (black, “exp”) are highlighted with a thicker line. The average Betti curves for *β*_1_-*β*_3_ of each computation of the surrogate data are drawn in thin color lines: uniform resampling of spike times (blue, “U”), exchange of spike times between collections (red, “EB”), exchange of spike times within collections (yellow, “EW”), Poisson generated spike data (purple, “P”). As in Fig. 2B, the two top (respectively, bottom) rows correspond to the increasing (resp., decreasing) filtration method, and the odd (resp., even) rows correspond to *k* = 0 (resp., *k* = 1).

To determine the consistency of these features across the 28 datasets, we used the integrated Betti values and the centers of mass of the Betti curves. Thus, for every fixed Betti number (*β*_1_, *β*_2_ or *β*_3_), we considered the difference between the integrated Betti value of each surrogate (mean value over the 20 computations) and the integrated Betti value of the experimental data, and averaged them over the 80 collections from each dataset (see Fig. 6 for *β*_1_ and Figs. S4-S5 for *β*_2_-*β*_3_). Consistent with Fig. 6, Figs. S4-S5, and with the single dataset of Fig. 5, the behavior of the experimental data departs from the behavior of the surrogates, especially for the mid-range values of *q* (5 to 50 sec^−1^). This holds both when the neuron of origin is ignored (*k* = 0) and when it is considered relevant (*k* = 1), and it is seen for both increasing and decreasing filtrations. All surrogates behave in a way that is distinct from the experimental data. As in Fig. 6, three of the surrogates (U, EB, EW) behave similarly to each other, while the Poisson surrogate (P) deviates from the data more extensively. We confirmed these observations using the centers of mass of the Betti curves in place of the integrated Betti values (see statistical tests below).

**Figure 6:**
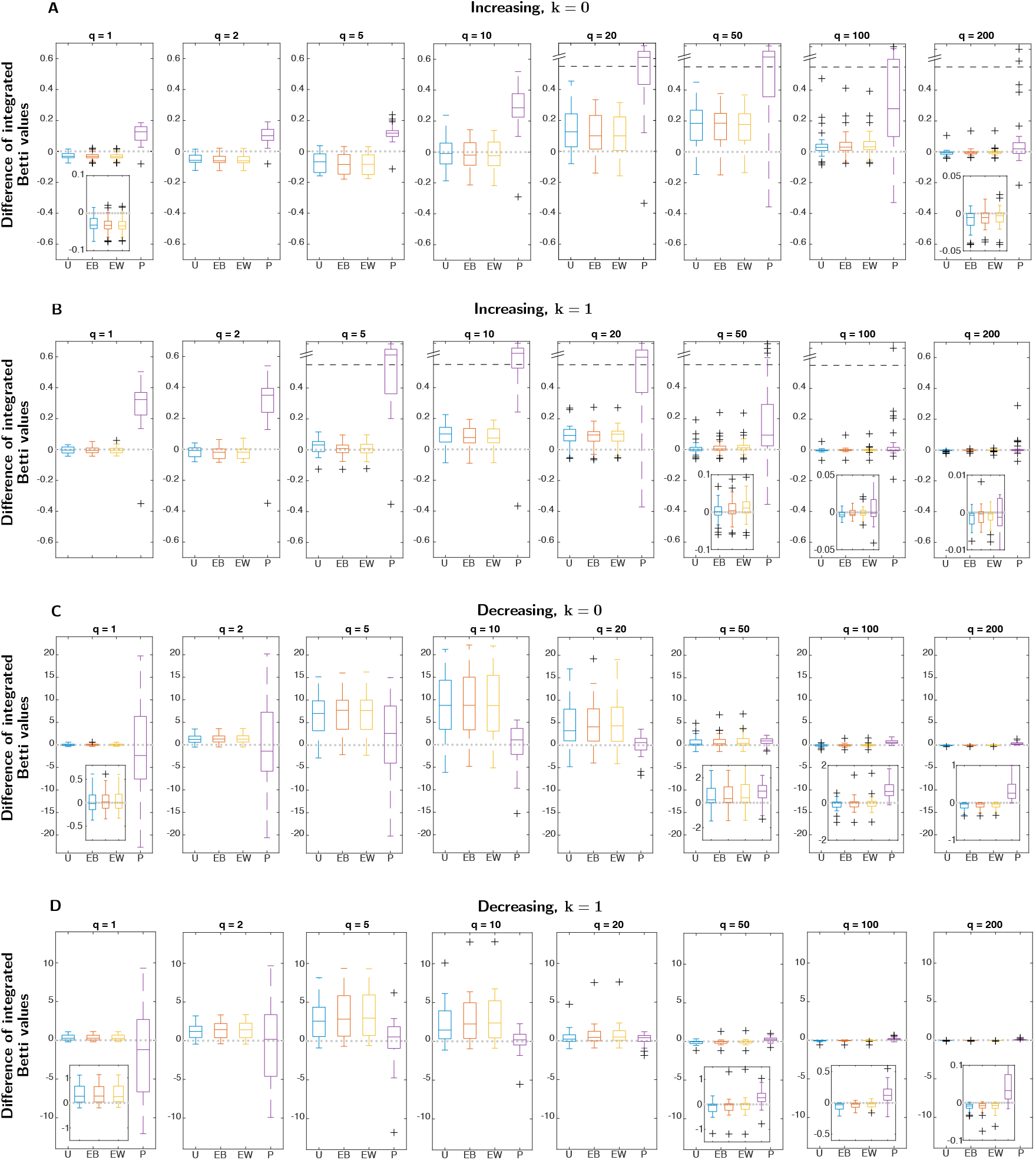
Distribution of the integrated Betti values for *β*_1_ of neural data and surrogates. Distribution across the 28 datasets of the difference between the integrated Betti values for *β*_1_ of each kind of surrogate, averaged over 20 examples, and the integrated Betti value of the experimental data. Each plot shows the four surrogates: uniform resampling of spike times (blue, “U”), exchange of spike times between collections (red, “EB”), exchange of spike times within collections (yellow, “EW”), Poisson generated spike data (purple, “P”). Insets zoom in on some boxplots at a smaller scale. When present, a dashed line bounds an area at the extreme of a plot beyond which data are shown on a compressed ordinate. The four panels are for increasing (**A**,**B**) and decreasing (**C**,**D**) filtrations, and for *k* = 0 (**A**,**C**) and *k* = 1 (**B**,**D**). For example, in **A**, the means of the integrated Betti values for the Poisson surrogates for all timescales, *q* are greater than the means of the integrated Betti values for the experimental spiking responses from the 28 datasets. Note that the deviation of the behavior of the surrogates is maximal for the range *q* = 5 sec ^−1^ to *q* = 50 sec^−1^.

Fig. 7 and Figs. S6-S7 assess the statistical significance of these observations for *β*_1_-*β*_3_ via the two-sample Kolmogorov-Smirnov (KS) test applied to the distributions of integrated Betti values. Even for the EW surrogate, which perturbs the recorded spike trains the least, the KS test statistic takes on large values, especially for mid-range values of *q*, rejecting (at the 0.05 significance level) the null hypothesis that the EW surrogate and the experimental data samples come from the same distribution. More generally, the analysis shows that the surrogates for U, EB, and EW diverge in a similar, timescale-dependent manner from the experimental data, while the Poisson generated data is always significantly different from both the experimental data and the other surrogates. These observations are confirmed by the KS test applied in the same way to the distributions of centers of mass of the Betti curves (Figs. S8-S10).

**Figure 7:**
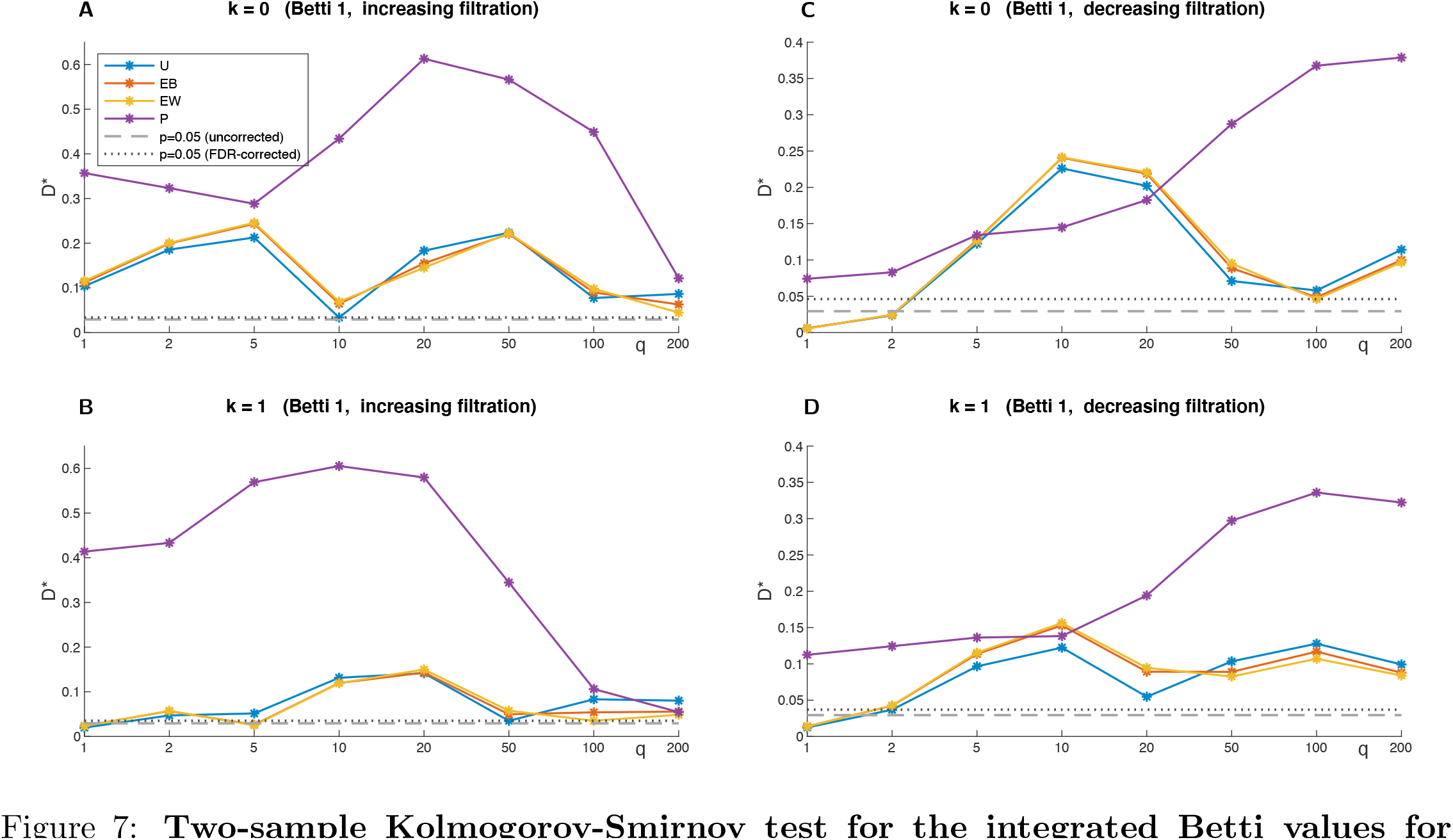
Two-sample Kolmogorov-Smirnov test for the integrated Betti values for *β*_1_. Two-sample Kolmogorov-Smirnov test statistic *D* ^*\**^(ordinate) for comparison of integrated Betti values for *β*_1_ of the experimental data with each type of surrogate data. The sample for the experimental data consists of the (28×80) integrated Betti values from all 80 collections from all 28 datasets. The samples for each surrogate consist of the (20 × 28 × 80) integrated Betti the samples of values of the 20 computations. Values above the dashed line, corresponding to *p* = 0.05, indicate rejection of the null hypothesis that the two samples come from the same distribution. The dotted line corresponds to the value *p* = 0.05 corrected for multiple comparisons, using the false discovery rate method (see Materials and Methods). The four panels are for increasing (**A**,**B**) and decreasing (**C**,**D**) filtrations, and for *k* =0 (**A**,**C**) and *k* = 1 (**B**,**D**).

In sum, Figs. 3-7 and S2-S10 demonstrate that the neural activity elicited by images with a range of statistical properties, and recorded in either V1 or V2, have a common topological structure that is distinct from that of surrogates with matching spike counts and spike timing distributions. These differences are present over a broad range of temporal scales, and are most prominent at a temporal scale of 5−50 sec^−1^, i.e., 20 to 200 ms.

The analysis so far shows that the observed firing patterns depart from that of surrogates that randomize spike times but maintain mean firing rates in various ways. This departure is manifest in the Betti curves, but the Betti curves are only an indirect reflection of the structure of the response space. To gain insight into this topological structure, we compared the measured Betti curves with those obtained from randomly-assigned distances or by sampling points in simple geometric spaces [22, 36, 50]. Specifically, we computed the Betti curves associated with: (i) a random symmetric 64×64 matrix with zeros on the diagonal [22], (ii) a sample of 64 random points in a unit (hyper)cube within a *d*-dimensional Euclidean space [22], and (iii) a sample of 64 random points in the hyperbolic ball model of the *d*-dimensional hyperbolic space [50]. For Euclidean and hyperbolic spaces, we considered dimensions *d* from 1 to 15. For the hyperbolic models, we also varied the effective curvature. As in [50], we implemented this by keeping intrinsic (Gaussian) curvature fixed at−1 but changing typical distances between the points (i.e., varying from 1 to 5 the maximum radius *R*_max_ of the hyperbolic ball model, see Materials and Methods). We therefore refer to models with low *R*_max_ as hyperbolic spaces with moderate curvature. For each random or geometric model, we computed the Betti curves of 300 sets of samples. We then compared these distributions with the Betti curves selected from our data via integrated Betti values and centers of mass. We restricted the analysis to collections with at most one empty response, in order to have 64 data points (and to exactly match the geometric models), and we randomly chose one such collection for every dataset that had collections meeting this criterion. This yielded Betti curves from 18 different datasets.

To determine compatibility with the geometric models, we first tallied how many of the experimental integrated Betti values were within 3 standard deviations of the mean of the 300 values of the model for all Betti numbers *β*_1_-*β*_3_ and for both increasing and decreasing filtrations. This procedure was followed for the grid of values of the *k* and *q* parameters of the Victor-Purpura distance (Fig. 8). We then compared the centers of mass using the same notion of compatibility

**Figure 8:**
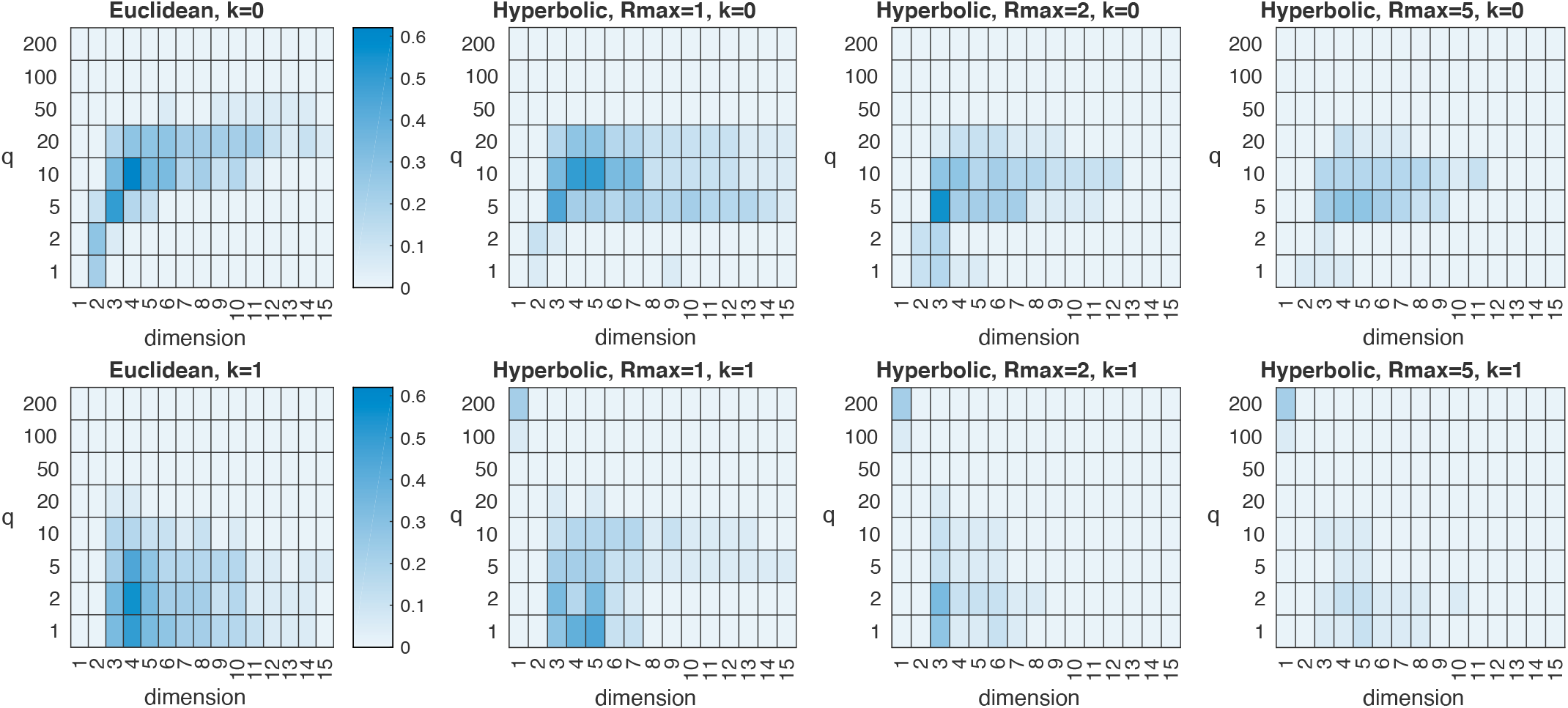
Compatibility of the experimental Betti curves with geometric models, based on integrated Betti values. The heatmaps show the fraction of Betti curves of 18 selected collections from different datasets which are compatible with the Euclidean (column 1) and hyperbolic models (*R*_max_=1, 2, 5) (columns 2-4) of dimension *d=*1, …, 15 (abscissas), for all values of the parameters *q* (ordinates) and *k* (rows) of the Victor-Purpura distance. The notion of compatibility we introduced requires the experimental integrated Betti values to be within 3 standard deviations of the mean of the 300 values of the model, for all Betti numbers *β*_1_-*β*_3_ and for both increasing and decreasing filtrations. In the range *q* = 5 to 20 sec^−1^, the greatest compatibility occurs for dimensions 3-5, and for the Euclidean or hyperbolic models with moderate curvature (*R*_max_ = 1). As we determined by visual inspection of the Betti curves, the hotspot at dimension 1 and *q* = 200 in the hyperbolic heatmaps for *k* = 1 is an artifact due to the fact that the low dimension constrains the Betti curves of the hyperbolic models to being close to (identically) zero, hence compatible in some cases with the experimental Betti curves at the extreme value *q* = 200.

(Fig. S11).

While none of these models were a good fit to the data, the Euclidean and low-curvature hyperbolic models (*R*_max_= 1) came the closest (Fig. 8). Fig. S11 shows however that compatibility is reduced when considering centers of mass instead of integrated Betti values, in particular for *k* = 1. Compatibility is maximal for low dimensions (*d* = 3 to 5) in the range we considered, consistently across the geometric models. In all cases, Betti curves associated with the experimental data were found to be incompatible with the random symmetric matrix model. Interestingly, across the geometric models, compatibility of integrated Betti values between the experimental data and the models (Fig. 8) is maximized in a consistent region of the parameter space of the Victor-Purpura distance. These regions correspond approximately to *q=*5 to 20 sec^−1^ for *k* = 0, and *q=*1 to 10 sec^−1^ for *k* = 1, while for large values of *q* the compatibility is very low in all cases.

## Discussion

### Geometry of spiking patterns

Understanding the properties of biological neural networks requires examining the structure of spike trains – the sequences and patterns of action potentials generated by groups of neurons in the cortex during the processing of information. We show here that characterizing the response space in which these patterns live through TDA provides insight into the intrinsic structure of the activity patterns generated in V1 and V2 in the macaque monkey during robust visual stimulation. The influence of timescale on this structure suggests that local networks in V1 and V2, and perhaps the interactions between these two cortical areas, play a major role in shaping the geometry of the spike response space, and in fact dominate the influence of the stimuli. While no simple model such as a circle, torus, or sphere [36] recapitulates the space, across a range of timescales the spaces extracted by TDA are most similar to low-dimensional models of Euclidean geometry or hyperbolic geometry with a modest amount of negative curvature (Figs. 8 and S11). Moreover, these spaces appear to be largely independent of the visual stimuli used in this study (Figs. 4 and S3) or the cortical region of the recordings (Figs. 3 and S2).

Note that while the spike pattern structures we identify are most consistent with Euclidean geometry or hyperbolic geometry with modest curvature, this geometry arises in a way that is fundamentally different from the manifolds and subspaces described above in the Introduction. The latter are derived from a vector state space of instantaneous spike rates [48], or calcium activity across a population [47], and the time evolution of these signals can be interpreted as trajectories across these subspaces. Here we begin with a family of metric state spaces, each defined by a timescale for comparing individual spike trains, and through TDA identify a family of spaces that each capture a unique spike pattern structure. These spike pattern structures exist in time as entities. Furthermore, vector space approaches assume that the actual timing of the spikes underlying the firing rates in the state space are largely irrelevant and can be effectively modeled as a Poisson point process, either homogeneous or inhomogeneous [36], with no temporal structure [25]. In our approach, we are specifically interested in the spike timing in individual and joint responses across groups of neurons.

Implicit in our results is the demonstration that TDA provides a way to characterize neuronal response spaces avoiding the limitations of vector space methods. TDA begins with the pairwise distances between instances of population activity. While several previous applications of TDA to neural spiking data have assumed a vector space structure or used a vector space embedding to compute these distances [22, 13, 14, 36], there is no fundamental reason to do so. Rather, the primary requirement is that the distance between two samples of neural activity corresponds to their dissimilarity. In [5], TDA methods combined with non-parametric (dis)similarity measures between spike trains were applied to synthetic spike train data for regimes classification in artificial neural networks. Here, we apply TDA in conjunction with a parametrized family of distances that have been shown to capture the meaningful dissimilarities between spike trains in several contexts [43, 1, 15, 19], and that are not Euclidean [4].

### Spiking responses driven by visual textures

Recognizing that neural activity is the result of an interaction of the inputs to the population and its intrinsic network properties, we sought to identify characteristics of the spiking response space that might apply to a wide range of visual stimuli. To this end, we used stimuli consisting of mathematically-defined visual textures [45, 49] of known relevance to natural vision [39]. Critically, this stimulus set included some textures whose visually-salient structure (first- and second-order spatial correlations) could be extracted by simple center-surround operations in the retina, and other textures whose structure (third- and fourth-order spatial correlations) can only be extracted by cortical computations, primarily in V2 [49]. As we show (Figs. 3-4 and S2-S3), the characteristics we identify are independent of this distinction; that is, these characteristics are shared by V1, where only first- and second-order spatial correlations are robust drivers of activity, and V2, where many neurons are sensitive to higher-order correlations [49]. During the data collection phase, we matched the orientation and spatial scale of the texture stimuli to the receptive field properties of the well-isolated single-units in the tetrode recordings. This allowed us to collect robust spiking activity even under propofol anesthesia/sufentanil analgesia [49]. These responses included strongly driven modulations of firing rate driven by many of the texture types, as well as trial-to-trial variability, thus facilitating wide exploration of the response space. We further promoted the exploration of the response space through the use of 64 distinct exemplars of each texture type. Finally, to ensure sensitivity to the details of spiking patterns, we based the analysis on individual spike trains rather than their averages, and used a distance that was sensitive to the timing of individual spikes rather than just overall spike count [36] or firing rate envelope [41].

### The importance of spiking pattern analysis in neuroscience

There is a considerable body of work spanning more than 3 decades, both experimental and theoretical, dedicated to elucidating the nature and function of spiking patterns in single neurons and across populations of neurons [18, 30, 9]. Several themes have emerged from this work. One is that the spiking patterns of individual neurons and groups of neurons can encode sensory information in the timing of their spikes [40, 43]. Second, the degree of synchrony of spiking patterns and the generation of discrete sequences of spikes across neurons, may play a role in transferring signals through synaptic pathways [33, 24]. Third, spiking patterns can arise through phase-locking of spikes to slower oscillatory signals in neural networks, a mechanism that is thought to mediate communication across cortical areas [28]. Fourth, spiking patterns in individual neurons may arise from joint activity in neural assemblies [35]. Finally, plasticity and learning in neural networks is dependent on the precise timing of pre- and post-synaptic spikes across the synapses in the network [8, 17]. Together, these themes emphasize the critical role spiking patterns play in brain function. However, the studies within this body of work use a wide range of approaches that of necessity often each provide only a partial picture of the spiking patterns of interest. Our study here is the first to apply TDA to the analysis of spiking patterns in the visual cortex. This study identifies a number of low-dimensional subspaces in the spiking response space, each at a different timescale. These subspaces reveal the presence of spiking patterns in the visual cortex generated during robust visual stimulation that do not encode the visual stimuli, but like the low-dimensional spaces that organize during spontaneous spiking activity [38], may reflect intrinsic states in visual cortical processing.

### Timescales and neurons of origin in spike patterns

Our approach recognizes that the timing of individual spikes is potentially important but it is agnostic as to what the relevant timescales are. This viewpoint is implemented by applying TDA to a family of distances controlled by a parameter *q* (in units of sec^−1^) that sets the resolution with which spike timing influences the measure of dissimilarity. Applying TDA to the resulting family of distances for *q=*1 to *q=*200 sec^−1^ allows us to focus on a range of timescales from 1sec down to 5ms.

The family of distances has a second parameter, *k*, which controls the importance of the neuron of origin in determining dissimilarity. For *k* = 0, the neuron of origin is irrelevant; for *k* = 1, changing the neuron of origin has unit cost. These extremes proved useful in analyzing multineuronal coding of spatial phase [3], demonstrating maximal information near *k* = 1; here, the dependence of response space geometry on *k* was relatively small.

Importantly, the distinction between experimental data and the random surrogates (U, EB, EW, P) depends systematically on the temporal resolution *q* of the distance between spike trains (Figs. 6-7 and S4-S10). The distinction is greatest for intermediate values of *q* = 5−50 sec^−1^, corresponding to resolutions of 200 to 20 ms). For low values of *q* (< 5), the distinction is generally lost, as the Victor-Purpura distance progressively disregards spike timing; for higher values (> 100), it is also generally weaker, indicating that at these timescales (with interspike intervals < 10ms) the systematic effect on geometry is less pronounced. Although the behavior of Poisson surrogates statistically diverge from the experimental data for the decreasing filtration method and high values of *q* (Figs. 5, 7 and S6-S10), we observe (Figs. 6-7 and S4-S10) that the distinction is clearly visible across the whole range of parameters and for both filtration methods. One can interpret the dependence between the timescale parameter *q* of the Victor-Purpura distance and the difference between experimental data and the random surrogates as the temporal scale of distinctive geometries that structure the spiking response space. It is notable that the timescales which most clearly reveal the geometry of the response space correspond to the timescales that are most informative for carrying visual information about contrast, spatial frequency, orientation, and texture – both as identified by analysis methods based on the Victor-Purpura distance [43], and by unrelated approaches [21]. Because the timescales at which the ongoing activity has the most distinctive geometric structure is similar to the timescales that are most informative about visual features, it is reasonable to hypothesize that this match facilitates transmitting visual information.

One way to interpret the results with the random surrogate data is to consider the degrees of freedom of the spiking activity. The Betti curves of the surrogates typically have higher values than the experimental data for intermediate values of *q* (Figs. 6 and S4-S5). This may indicate that the response space for the real spiking activity is more constrained than the synthetic spiking activity. In this range (*q=*5−50 sec^−1^), the random manipulation of the spike data in different ways consistently opens up holes in the response space, leading to higher Betti values (Figs. 5-6 and S4-S5). This result suggests that the neural response space produced during visual stimulation has fewer discontinuities (is more structured) than the response spaces produced by the surrogates.

### Comparison with simple geometric models

The comparison with random and geometric models highlights complementary aspects of the geometry of the space of sampled spike trains endowed with Victor-Purpura distances. Upon defining a notion of compatibility that accounts for the Betti curves of *β*_1_-*β*_3_ and both increasing and decreasing filtration, we observed that even if none of the considered models consistently fits the data, Euclidean geometry and hyperbolic geometry with moderate curvature (*R*_max_ =1) are more compatible with the geometry of the spike train responses (Figs. 8 and S11). Compatibility is concentrated in low dimensions of the geometric models (*d=*3-5). This observation is consistent with other geometrical descriptions of neural population activity [20]. In these descriptions, the instantaneous firing rates of a large number of neurons are found to occupy a low-dimensional manifold within a high-dimensional spiking response space. While in the study described here we analyze recordings from at most four neurons, the distances we consider between spike trains are intrinsic and do not depend on the choice of an embedding, and as a consequence, potentially capture aspects of the neural activity manifold. However, recordings of larger numbers of neurons are needed to demonstrate that this is the case. Fig. 8 also shows that the compatibility between the data and the geometric models assessed via integrated Betti values systematically depends on the timescale *q* of the Victor-Purpura distance between spike trains, and is maximized for mid-range values *q=*5−20 sec^−1^ when the neuron of origin of each spike is disregarded (*k* = 0), and for low-range values *q* = 1 − 5 sec ^−1^ when the origin of each spike is accounted for (*k* = 1). When assessed via centers of mass, however, compatibility is reduced for *k* = 0 and very low for *k* = 1 (Fig. S11), even if a systematic dependence on *q* is still observable.

### Considerations regarding the experimental procedure

The data analyzed here were collected from monkeys under propofol anesthesia/sufentanil analgesia and neuromuscular blockade. As a consequence, fixational eye movements could not have contributed to the fluctuations in spiking activity we observed in V1 and V2, but conversely, the natural dynamics of the visual input due to such eye movements is only partially approximated by the transient mode of stimulus presentation that we used. Another caveat is that the anesthesia and opiate analgesia may have produced noise correlations in the neural activity that contributed to the topological structure we extracted through TDA. While it is impossible to rule out any impact of anesthesia and analgesia on our results, there are two reasons that it is probably minor. First, as mentioned above, the timescales associated with the most consistent and distinctive state-space structure observed here corresponded to the timescales that are most important for carrying visual information in neurons in the visual cortex of the awake macaque [43]. Second, available evidence suggests that the effect of drug-induced changes in network state would be more likely to dilute any underlying structure, than to create it. Specifically, Ecker and co-workers [16] compared the variability of the spiking activity in V1 in awake monkeys with that seen in monkeys under sufentanil, inferring the presence of a sufentanil-related state variable. This state variable fluctuated with a timecourse that varied from 50 to 1650ms. If similar sufentanil-related fluctuations were present during our recording sessions, the impact would have been distributed randomly across the many 320ms spiking response samples.

### Methodological innovations

In summary, we introduce a new framework for applying TDA to spike train data, and use it to analyze patterns of spiking response in macaque visual cortex. There are two main methodological innovations. First, in contrast to most previous applications of TDA to neural data, we do not assume that the spike trains have a vector space embedding that induces distances and correlation measures between them; rather, the topological analysis is directly applied to dissimilarity measures (e.g. distances) that result from considering spike trains to be sequences of events – in this case, the Victor-Purpura spike train distances, which are non-Euclidean. A second innovation of the approach is the filtration – the sequence of simplicial complexes derived from the dissimilarities that are used to compute the Betti curves. Typically, an increasing filtration is used for TDA [37, 10]: graphs are progressively filled in for pairs of points at greater and greater dissimilarities; here we show that the decreasing filtration can reveal a clearer picture of the data’s geometry and topology. Betti curves, especially when averaged over several computations, capture the statistical distribution of the persistent topological features (tunnels, voids, etc.) across a filtration, which serves as a topological descriptor of the underlying metric spaces. In contrast to the usual filtration of graphs by increasing weights and their associated clique complexes, the decreasing filtration is obtained by considering, in reverse order, the *independence complex* of each graph (i.e., the clique complex of the complement of the graph). Betti curves of low dimension (*β*_1_–*β*_3_) of the decreasing filtration therefore carry information on the arrangements of high-order cliques of the original graphs, which is not related to the homology of the increasing filtration. Because the analysis is carried out for a sequence of distances parameterized by timescale, we are able to identify the range in which the geometric structure of the spiking response space is most distinctive: the range 5−50 sec^−1^, i.e., 20 to 200ms. As noted above, this timescale corresponds to the temporal precision that is most informative for decoding visual information from spike trains. This matching of the timescale of response space geometry and the timescale of neural coding may be a general feature of brain networks, and we speculate that networks in other domains (e.g., motor planning, learning, decision-making, etc.) will behave similarly.

## Materials and Methods

### Experiments

All procedures followed the guidelines provided by the US National Institutes of Health and Weill Cornell Medical College Animal Care and Use Committee. Full details concerning the physiological preparation and multi-tetrode single-unit recordings can be found in [34] and [49]. Detailed descriptions of the visual stimuli, their generation, and their display during the experimental sessions are given in [49], which also details how single-unit activity was characterized, and how the time-series of neural firing events, including the procedures utilized for spike sorting, were extracted from the multi-tetrode recordings. These methods are summarized in SI Materials and Methods.

### Visual stimuli and stimulation protocols

The visual stimuli used in this study are 16×16 black and white checkerboard-like patterns drawn from a mathematically-defined stimulus space. The stimulus space has 10 coordinate axes specifying the type of the multipoint correlations in each 2×2 grid of “checks” in the patterns, and the coordinates along these axes define the strength of the correlation (the fidelity with which the multipoint correlation is rendered across each pattern), and the correlation’s polarity [45, 49]. The 10 coordinate axes can be partitioned into four classes according to the order of the multipoints correlations [45]: first-, second-, third- and fourth-order correlations (see Fig. 1). In this study, the patterns were drawn from four coordinate values (2 magnitudes × 2 polarities) along each of the 10 axes in the stimulus space. The size of the 16 × 16 patterns, their position on the visual display screen, the orientation of the patterns, and the check-sizes in the patterns were chosen to optimize stimulation of isolated clusters of neurons based on the neurons’ receptive field properties as determined by the responses to sinusoidal gratings. Further details on this point are given in [49]. The topological data analysis presented here was performed on a subset of the neural responses obtained during the experimental runs. The experimental runs used (at least) 64 samples from the 4 coordinate values along each of the 10 axes in the stimulus space. In addition, 64 examples of fully random patterns were included in the experimental runs. The topological data analysis excluded the responses to the random patterns. For the experimental runs, a total of 2624 unique stimuli (64×41) each repeated 4 times (for a grand total of 10496 stimuli) was shown. In 20 of the 28 analyzed datasets, the 64 analyzed responses to visual stimuli of the same type are part of a larger sequence of 128 responses to 128 unique stimuli; in these cases, we selected for our analysis the responses to the first 64 unique stimuli shown during the experiment. The patterns were presented in extended pseudorandom sequences at 100% contrast, surrounded by a mid-level gray background. Each pattern appeared for 0.32 seconds and was immediately followed by the next pattern in the sequence. A different pseudorandom sequence was used for each of the four times a sequence was run during a recording block. There was a pause of about 1 minute between these repeats, during which time the gray background was displayed.

### Analyzed data

For each of the 28 datasets, the responses to all non-random stimulus types (each determined by a choice of an axis and a strength level for the correlations) for all repeats were then assembled to yield a total of 40×4 =160 collections. To study the topology of the space that neural activity occupies, we selected the 80 collections in each dataset with the smallest number of empty responses (i.e., responses with 0 spikes). The vast majority of these collections had at least 60 non-empty responses; see Fig. S1 (D). This choice for selecting collections is motivated by the fact that our analysis maps spike trains with at least one spike to distinct points in the topological space, while all empty spike trains are mapped to the same point. Thus, this selection criterion maximized the sampling of the space occupied by a collection.

### Data analysis and TDA

#### Victor-Purpura distance

The (multineuron) Victor-Purpura distance [43] is a cost-based distance between spike trains that has two parameters: a parameter *q* that controls the timescale used to quantify dissimilarity in spike timing, and a parameter *k* that determines the relevance of the neuron of origin of each spike. The distance between two neuronal responses is defined as the minimum cost of transforming one spike train into the other via a sequence of basic moves: addition or deletion of a single spike, with a cost of 1, shift of a single spike by a time interval Δ*t*, with a cost *q*|Δ*t*|, or change in the neuron of origin, with cost *k*. Thus, *q=*0 corresponds to ignoring spike time but retaining spike count, *q* > 0 corresponds to considering temporal structure at a scale of 1/ *q*. If a spike time in one train is within 1/ *q* of a spike time in a second train, they are considered to correspond, as they contribute less than 1 unit to the dissimilarity. Conversely, if spikes are more than 2 /*q* apart, they are considered to be unrelated, since shifting them into correspondence would incur a higher cost than deleting the spike from one train and inserting it into the other. Similarly, while for *k* = 0 changing the neuron of origin of a spike comes with no cost, for *k* = 1 the corresponding cost is equivalent to the insertion of a spike. Further background on the Victor-Purpura distance, as well as efficient dynamic-programming algorithms for calculating it, can be found in [44, 2]. In our analysis, for each collection of neuronal responses we determined the distance matrix *D=*(*D*_*ij*_), where *D*_*ij*_ is the Victor-Purpura distance with parameters (*q, k)*between the *i*th and the *j*th response in the collection. The parameters *q* and *k* ranged over a grid of values: *q=*1, 2, 5, 10, 20, 50, 100, 200 (sec^−1^), and *k* = 0, 1. The Victor-Purpura distance between spike trains is computed using the “labdist faster qkpara opt” Matlab function implemented by Thomas Kreuz, available at: http://www-users.med.cornell.edu/~jdvicto/labdist_faster_qkpara_opt.html

### TDA methods

We applied and extended the *clique topology* method introduced in [22], which corresponds to computing the Betti curves associated with our increasing filtration. Given a symmetric matrix *M* with zeros on its main diagonal, the first step of the method consists in considering the above-diagonal part of the matrix, transforming it by rank ordering its entries (thus replacing the original entry values by natural numbers 0, 1, …), and completing the below-diagonal part of the rank-ordered matrix by symmetry. The method then computes the persistent homology (see TDA software) of the rank-ordered matrix and finally determines the associated Betti curves. Since the input matrix is transformed by considering only the rank ordering of its entries, the output is invariant to monotonic transformation applied entry-wise to *M*. Given a dissimilarity or distance matrix, the clique topology method orders the entries increasingly. We observe that in [22] the method is also applied to a matrix *C* = (*C*_*ij*_) of correlations by transforming it into a dissimilarity matrix *D=(D*_*ij*_)via the application of any function that inverts the ordering between the absolute values l*C*_*ij*_l of the correlations and the entries *D*_*ij*_, e.g., *D*_*ij*_= −l*C*_*ij*_l. In this work, we apply the clique topology method by ordering the entries of our distance matrices both increasingly (*increasing filtration*) and decreasingly (*decreasing filtration*), the latter case corresponding to the *weight rank clique* filtration method introduced in [32].

A sequence of nested simple graphs *G*_*0*_⊂ *G*_*1*_⊂ … ⊂ *G*_*s*_, called a filtration, can be determined from a rank-ordered *n*×*n* symmetric matrix (typically representing pairwise distances) as follows. Each graph has the same set of *n* nodes, and the edges of a graph *G*_*k*_ are determined by thresholding the rank-ordered matrix (setting to one all entries smaller than *k*, and setting to zero the remaining entries) and regarding it as an adjacency matrix. An edge is therefore present between nodes *i* and *j* of the graph *G*_*k*_ if and only if the (*i, j)*entry of the rank-ordered matrix is less than *k*. If the above-diagonal entries of the symmetric rank-ordered matrix are all different, the graph *G*_0_ has no edges, the graph *G*_1_ has one edge, corresponding to the smallest off-diagonal entry of the rank-ordered matrix, and so on. As the edges fill in, graphs can be enriched with higherorder connectivity information encoded by basic “pieces” of different dimensions, called simplices. Specifically, a *p-clique* (subgraph of *p* all-to-all connected nodes) in a graph is regarded as a (*p*−1) - dimensional simplex: 2-cliques are the edges of the graph and are viewed as line segments (1-dimensional simplices), 3-cliques are viewed as triangles (2-dimensional simplices), 4-cliques are viewed as tetrahedra (3-dimensional simplices), etc. In this way, each graph becomes a *simplicial complex*, called the *clique complex* of the graph, in which arrangements of cliques can enclose “holes” of different dimensions. The number of 1-dimensional tunnels, 2-dimensional voids and 3dimensional “cavities”, known as Betti numbers and denoted by *β*_1_, *β*_2_ and *β*_3_ respectively, are computed at each step of the filtration. The Betti numbers are viewed as functions of the edge density *ρ*, the number of filled-in edges divided by the number *N* = *n*(*n* − 1)/2 of potential edges. These functions (*β*_1_(*ρ*), *β*_2_(*ρ*) and *β*_3_(*ρ*)) are the Betti curves, which we computed over the range from *ρ* = 0 to 0.6 in our analysis.

### TDA software

To compute clique topology and persistent homology, we use a faster and equivalent alternative to the original CliqueTop scripts (https://github.com/nebneuron/clique-top; see [22]), namely Ripser [6] (https://github.com/Ripser/ripser) on a rank-ordered distance matrix. Starting from a distance matrix *D=(D*_*ij*_), we added small (symmetric) Gaussian noise and transformed the resulting symmetric matrix by rank-ordering its entries in increasing (resp., decreasing) order for the increasing (resp., decreasing) filtration method. Note that the purpose of the Gaussian noise is to uniquely specify an ordering of the entries in presence of equal values, and its magnitude was chosen small enough to preserve the ordering between entries with different values. Ripser computes the persistent homology of the clique filtration associated with the ordered matrix and outputs the so-called *barcode*, a collection of pairs of real numbers *B={(b*_*i*_, *d*_*i*_)}_*i*=1,…,*m*_ for the chosen dimension of homology, which are *j* = 1, 2, 3 in our setting. The pairs (*b*_*i*_, *d*_*i*_) for which a particular void of dimension *j* is present. If the size of the input matrix is *n* × *n*, the Betti curve is obtained from the barcode *B* by setting *β*_*j*_ (*r*/*N*) to the number of (*b*_*i*_, *d*_*i*_) in *B* such are the ranges in edge density that *b*_*i*_≤ *r* < *d*_*i*_, where *N* = *n*(*n* − 1)/2 and *r* is any integer between 0 and the maximum value *s* such that *s*/*N* does not exceed a maximum edge density *ρ*_max_. The parameter *ρ*_max_ was set to 0.6 in our analysis for computational efficiency, following [22]. The function *β*_*j*_(*ρ*) for *ρ* ∈ [0, *ρ*_max_] is therefore piecewise-constant, as it is constant on every interval [*r*/*N*, (*r* + 1)/*N*).

### Integrated Betti values and centers of mass

In the analysis we use integrated Betti values [22] (see Figs. 3-4, 6-8 and S4-S7) and centers of mass (see Figs. S2-S3 and S8-S11) to summarize the shape of Betti curves. Let *β:* [0, *ρ*_max_]→ ℝ be a Betti curve. Its integrated Betti value is defined as

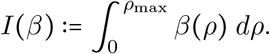

The center of mass of the Betti curve is defined as

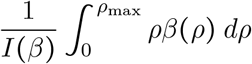

when *I*(*β*)>0, and it is defined to be zero when *I*(*β*) = 0.

### Boxplots and KS statistical test

Boxplots are generated with the Matlab function “boxplot”. The two-sample Kolmogorov-Smirnov (KS) test was performed using the Matlab function “kstest2”. In Fig. 7 and Figs. S6-S10 the significance level of the KS test has been corrected for multiple comparisons via the false discovery rate correction [7].

### Surrogate spike train data

In our analysis, we compared the Betti curves computed from the experimental data to the Betti curves of four types of surrogate spike train data (Figs. 5-7 and S4-S10). For each experimental dataset, we generated 20 surrogate spike datasets for each of the four types. The four types of surrogate spike train data were generated by perturbing specific aspects of the spiking patterns in the original experimental data.

1. *Uniform resampling of spike times* (U). Each sequence of spikes from each neuron from each response is replaced by a sequence of the same number of spikes randomly distributed over the length of the response interval (0−320ms). The sequence of labels indicating which neuron fired the spikes is preserved, as is the number of spikes of each neuron in the response.
2. *Exchange resampling of spike times between collections* (EB). Spikes are randomly swapped between the 80 selected collections of a dataset, preserving their time of occurrence in the 320ms response and their neuron of origin. Thus, each response in the surrogate dataset has the same number of spikes as the original, and the overall distribution of spike times across the dataset are maintained.
3. *Exchange resampling of spike times within collections* (EW). As in 2, but with the swapping restricted to responses within each collection. Thus, spike counts are unchanged within each response, as is the distribution of spike times within each collection.
4. *Poisson generated spike trains* (P). For each neuron contributing to a dataset, we determined its overall firing rate across the dataset. We then replaced each neuron’s spike train in all non-empty responses in the dataset with a sample of a Poisson process with that rate. We set the bin width in the Poisson generator to 0.0001ms. Note that this surrogate dataset does not preserve the number of spikes of each neuron in each response, only the average.

### Geometric models

Models of random and geometric spaces were considered for Fig. 8 and Fig. S11. The random symmetric matrices and the distance matrices of random points in a Euclidean space were generated following [22], while the distance matrices of random points in a hyperbolic space were generated following [50].

#### 1. Random symmetric matrix model

The nonzero (off-diagonal) elements of the distance matrix *D=(D*_*ij*_)are randomly-chosen positive numbers uniformly distributed in the interval 0, 1, with *D*_*ij*_ = *D*_*ji*_, unconstrained by the triangle inequality.

#### 2. Euclidean geometry model

To generate a random Euclidean distance matrix *D D*_*ij*_, 64 random points are uniformly sampled in a unit (hyper)cube within the *d*-dimensional Euclidean space, for a fixed dimension *d* between 1 and 15. Each entry *D*_*ij*_ is set equal to the Euclidean distance between the randomly-chosen *i*th and *j*th points.

#### 3. Hyperbolic geometry model

Similarly to [50], we generated a random hyperbolic distance matrix *D* = (*D*_*ij*_) by uniformly sampling 64 points in the *d*-dimensional hyperbolic space (for a fixed dimension *d* between 1 and 15), using the hyperbolic ball model [29] with curvature *ζ=*1. We sampled a standard *d*-variate Gaussian (using the Matlab function “randn”) and rescaled the radii of the sampled points by selecting radii *r* within [0, *R*_max_] following the distribution *ρ*(*r*)∼ sinh((*d* − 1)*r*). We examined *R*_max_ values in the range from 1 to 10, and results for *R*_max_ = 1, 2, 5 are shown in Fig. 8 and Fig. S11. The distance *D*_*ij*_ between two points of radii *r*_*i*_ and *r*_*j*_ with an angle Δ*θ* between them is determined from the hyperbolic law of cosines

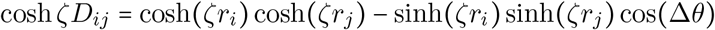

with curvature *ζ=*1 in our setting.

### Data and code availability

The data related to this article is available at https://github.com/aguidolin/visual-spike. A set of Matlab functions and scripts to replicate the analysis is available at https://github.com/aguidolin/topspike. Additional code related to this article may be requested from the authors.

## Acknowledgments

JDV and KPP acknowledge Qin Hu, Ferenc Mechler, Eyal Nitzany, Anita Schmid, Dan Thengone and Yunguo Yu for assisting with data collection. They also acknowledge NIH grant numbers EY09314 and EY07977 from the National Eye Institue of the NIH. SR was supported by Ikerbasque (The Basque Foundation for Science); AG and SR were supported by the Basque Government through the BERC 2018-2021 program and by the Ministry of Science, Innovation and Universities: BCAM Severo Ochoa accreditation SEV-2017-0718 and through project RTI2018-093860B-C21 funded by (AEI/FEDER, UE) with acronym “MathNEURO”. AG acknowledges the support of the Wallenberg AI, Autonomous Systems and Software Program (WASP) funded by the Knut and Alice Wallenberg Foundation. MD and SR acknowledge suppport of Inria through the Associated Team “NeuroTransfSF”.

## Additional information

### Author contribution

All authors designed the research; KPP and JDV carried out the physiological experiments; AG carried out the computational analyses; SR and JDV provided funding; AG, JDV, KPP, and SR wrote the initial draft; all authors contributed to the final draft.

## References

[1] Alyssa Accomando, Carlos Vargas-Irwin, and James Simmons. Neural spike train similarity algorithm detects differences in temporal patterning of bat echolocation call sequences. In Proceedings of Meetings on Acoustics 173EAA, volume 30, page 010002. Acoustical Society of America, 2017.

[2] Dmitriy Aronov. Fast algorithm for the metric-space analysis of simultaneous responses of multiple single neurons. Journal of neuroscience methods, 124(2):175–179, 2003.

[3] Dmitriy Aronov, Daniel S Reich, Ferenc Mechler, and Jonathan D Victor. Neural coding of spatial phase in V1 of the macaque monkey. Journal of Neurophysiology, 89(6):3304–3327, 2003.

[4] Dmitriy Aronov and Jonathan D Victor. Non-Euclidean properties of spike train metric spaces. Physical Review E, 69(6):061905, 2004.

[5] Jean-Baptiste Bardin, Gard Spreemann, and Kathryn Hess. Topological exploration of artificial neuronal network dynamics. Network Neuroscience, 3(3):725–743, 2019.

[6] Ulrich Bauer. Ripser: efficient computation of Vietoris–Rips persistence barcodes. Journal of Applied and Computational Topology, pages 1–33, 2021.

[7] Yoav Benjamini and Yosef Hochberg. Controlling the false discovery rate: a practical and powerful approach to multiple testing. Journal of the Royal statistical society: series B (Methodological), 57(1):289–300, 1995.

[8] Guo-qiang Bi and Mu-ming Poo. Synaptic modifications in cultured hippocampal neurons: dependence on spike timing, synaptic strength, and postsynaptic cell type. Journal of neuroscience, 18(24):10464–10472, 1998.

[9] Peter Cariani and Janet M Baker. Time is of the essence: Neural codes, synchronies, oscillations, architectures. Frontiers in Computational Neuroscience, 16, 2022.

[10] Gunnar Carlsson. Topology and data. Bulletin of the American Mathematical Society, 46(2):255–308, 2009.

[11] SueYeon Chung and LF Abbott. Neural population geometry: An approach for understanding biological and artificial neural networks. Current opinion in neurobiology, 70:137–144, 2021.

[12] John P Cunningham and M Yu Byron. Dimensionality reduction for large-scale neural recordings. Nature Neuroscience, 17(11):1500–1509, 2014.

[13] Carina Curto and Vladimir Itskov. Cell groups reveal structure of stimulus space. PLoS Computational Biology, 4(10):e1000205, 2008.

[14] Yuri Dabaghian, Facundo Mémoli, Loren Frank, and Gunnar Carlsson. A topological paradigm for hippocampal spatial map formation using persistent homology. PLoS Computational Biology, 2012.

[15] Patricia M Di Lorenzo and Jonathan D Victor. Taste response variability and temporal coding in the nucleus of the solitary tract of the rat. Journal of Neurophysiology, 90(3):1418–1431, 2003.

[16] Alexander S Ecker, Philipp Berens, R James Cotton, Manivannan Subramaniyan, George H Denfield, Cathryn R Cadwell, Stelios M Smirnakis, Matthias Bethge, and Andreas S Tolias. State dependence of noise correlations in macaque primary visual cortex. Neuron, 82(1):235–248, 2014.

[17] Daniel E Feldman. The spike-timing dependence of plasticity. Neuron, 75(4):556–571, 2012.

[18] Jean-Marc Fellous, Paul HE Tiesinga, Peter J Thomas, and Terrence J Sejnowski. Discovering spike patterns in neuronal responses. Journal of Neuroscience, 24(12):2989–3001, 2004.

[19] Luca A Finelli, Seth Haney, Maxim Bazhenov, Mark Stopfer, and Terrence J Sejnowski. Synaptic learning rules and sparse coding in a model sensory system. PLoS Computational Biology, 4(4):e1000062, 2008.

[20] Peiran Gao and Surya Ganguli. On simplicity and complexity in the brave new world of large-scale neuroscience. Current opinion in neurobiology, 32:148–155, 2015.

[21] Timothy J Gawne and Barry J Richmond. How independent are the messages carried by adjacent inferior temporal cortical neurons? Journal of Neuroscience, 13(7):2758–2771, 1993.

[22] Chad Giusti, Eva Pastalkova, Carina Curto, and Vladimir Itskov. Clique topology reveals intrinsic geometric structure in neural correlations. Proceedings of the National Academy of Sciences, 112(44):13455–13460, 2015.

[23] Tijl Grootswagers, Amanda K Robinson, Sophia M Shatek, and Thomas A Carlson. Untangling featural and conceptual object representations. NeuroImage, 202:116083, 2019.

[24] Richard HR Hahnloser, Alexay A Kozhevnikov, and Michale S Fee. An ultra-sparse code underlies the generation of neural sequences in a songbird. Nature, 419(6902):65–70, 2002.

[25] Olivier J Hénaff, Yoon Bai, Julie A Charlton, Ian Nauhaus, Eero P Simoncelli, and Robbe LT Goris. Primary visual cortex straightens natural video trajectories. Nature communications, 12(1):1–12, 2021.

[26] Olivier J Hénaff, Robbe LT Goris, and Eero P Simoncelli. Perceptual straightening of natural videos. Nature neuroscience, 22(6):984–991, 2019.

[27] Ann M Hermundstad, John J Briguglio, Mary M Conte, Jonathan D Victor, Vijay Balasubramanian, and Gašper Tkačik. Variance predicts salience in central sensory processing. eLife, 3:e03722, 2014.

[28] Adam Kohn, Anna I Jasper, Joao D Semedo, Evren Gokcen, Christian K Machens, and M Yu Byron. Principles of corticocortical communication: proposed schemes and design considerations. Trends in Neurosciences, 43(9):725–737, 2020.

[29] Dmitri Krioukov, Fragkiskos Papadopoulos, Maksim Kitsak, Amin Vahdat, and Marian Boguná. Hyperbolic geometry of complex networks. Physical Review E, 82(3):036106, 2010.

[30] Artur Luczak, Bruce L McNaughton, and Kenneth D Harris. Packet-based communication in the cortex. Nature Reviews Neuroscience, 16(12):745–755, 2015.

[31] Marino Pagan, Luke S Urban, Margot P Wohl, and Nicole C Rust. Signals in inferotemporal and perirhinal cortex suggest an untangling of visual target information. Nature neuroscience, 16(8):1132–1139, 2013.

[32] Giovanni Petri, Martina Scolamiero, Irene Donato, and Francesco Vaccarino. Topological strata of weighted complex networks. PLoS one, 8(6):e66506, 2013.

[33] Alex D Reyes. Synchrony-dependent propagation of firing rate in iteratively constructed networks in vitro. Nature neuroscience, 6(6):593–599, 2003.

[34] Anita M Schmid, Keith P Purpura, and Jonathan D Victor. Responses to orientation discontinuities in V1 and V2: physiological dissociations and functional implications. Journal of Neuroscience, 34(10):3559–3578, 2014.

[35] Hideaki Shimazaki, Shun-ichi Amari, Emery N Brown, and Sonja Grün. State-space analysis of time-varying higher-order spike correlation for multiple neural spike train data. PLoS computational biology, 8(3):e1002385, 2012.

[36] Gurjeet Singh, Facundo Memoli, Tigran Ishkhanov, Guillermo Sapiro, Gunnar Carlsson, and Dario L Ringach. Topological analysis of population activity in visual cortex. Journal of vision, 8(8):11–11, 2008.

[37] Ann E Sizemore, Jennifer E Phillips-Cremins, Robert Ghrist, and Danielle S Bassett. The importance of the whole: topological data analysis for the network neuroscientist. Network Neuroscience, 3(3):656–673, 2019.

[38] Carsen Stringer, Marius Pachitariu, Nicholas Steinmetz, Charu Bai Reddy, Matteo Carandini, and Kenneth D Harris. Spontaneous behaviors drive multidimensional, brainwide activity. Science, 364(6437):eaav7893, 2019.

[39] Tiberiu Tesileanu, Mary M Conte, John J Briguglio, Ann M Hermundstad, Jonathan D Victor, and Vijay Balasubramanian. Efficient coding of natural scene statistics predicts discrimination thresholds for grayscale textures. eLife, 9:e54347, 2020.

[40] Frédéric Theunissen and John P Miller. Temporal encoding in nervous systems: a rigorous definition. Journal of computational neuroscience, 2(2):149–162, 1995.

[41] Jonathan D Victor. Spike train metrics. Current Opinion in Neurobiology, 15(5):585–592, 2005.

[42] Jonathan D Victor and Mary M Conte. Local image statistics: maximum-entropy constructions and perceptual salience. JOSA A, 29(7):1313–1345, 2012.

[43] Jonathan D Victor and Keith P Purpura. Nature and precision of temporal coding in visual cortex: a metric-space analysis. Journal of Neurophysiology, 76(2):1310–1326, 1996.

[44] Jonathan D Victor and Keith P Purpura. Metric-space analysis of spike trains: theory, algorithms and application. Network: Computation in Neural Systems, 8(2):127–164, 1997.

[45] Jonathan D Victor, Daniel J Thengone, Syed M Rizvi, and Mary M Conte. A perceptual space of local image statistics. Vision research, 117:117–135, 2015.

[46] Ryan C Williamson, Benjamin R Cowley, Ashok Litwin-Kumar, Brent Doiron, Adam Kohn, Matthew A Smith, and Byron M Yu. Scaling properties of dimensionality reduction for neural populations and network models. PLoS Computational Biology, 12(12):e1005141, 2016.

[47] Yang Xie, Peiyao Hu, Junru Li, Jingwen Chen, Weibin Song, Xiao-Jing Wang, Tianming Yang, Stanislas Dehaene, Shiming Tang, Bin Min, and Liping Wang. Geometry of sequence working memory in macaque prefrontal cortex. Science, 375(6581):632–639, 2022.

[48] Byron M Yu, John P Cunningham, Gopal Santhanam, Stephen I Ryu, Krishna V Shenoy, and Maneesh Sahani. Gaussian-process factor analysis for low-dimensional single-trial analysis of neural population activity. Journal of Neurophysiology, 102(1):614–635, 2009.

[49] Yunguo Yu, Anita M Schmid, and Jonathan D Victor. Visual processing of informative multipoint correlations arises primarily in V2. eLife, 4:e06604, 2015.

[50] Yuansheng Zhou, Brian H Smith, and Tatyana O Sharpee. Hyperbolic geometry of the olfactory space. Science Advances, 4(8):eaaq1458, 2018.

