## Supplementary Information for "Geometry of spiking patterns in early visual cortex: a Topological Data Analytic approach"

September 12, 2022

#### Materials and methods

##### Physiologic methods

Standard techniques [1, 3], consistent with US National Institutes of Health and Weill Cornell Medical College Animal Care and Use Committee guidelines, were used to prepare six adult macaque monkeys for acute extracellular recordings from areas V1 and V2. After initial sedation with ketamine and isoflurane, general anesthesia was induced and maintained with infusion of a propofol/sufentanil mixture during surgical preparation and recording. All incision sites were infiltrated with bupivacaine prior to tissue opening. Endotracheal tube placement and catheterization of one femoral artery, both femoral veins, and the urethra, were completed after initial sedation. Hydration was provided with Normosol and dextrose, and eye movements were minimized by neuromuscular blockage through infusion of vecuronium or rocuronium bromide. Heart rate and rhythm, arterial blood pressure, body temperature, end-tidal CO2 partial pressure, arterial oxygen saturation, EEG, and urine output were monitored over the entire course of each experiment. The pupils were dilated with topical atropine and the eyes were protected with gas-permeable contact lenses and topical flurbiprophen. Animal maintenance also included administration of penicillin and dexamethasone at the start of the experiment and gentamicin later in the recording session, if needed. All combined, the measures briefly described here maintained the animals in a stable physiological state for 4-5d.

##### Recording and recovery of recording sites

Following a craniotomy, a small dural incision was made and three to six guide tubes, each containing a quartz-platinum-tungsten tetrode (Thomas Recording GmbH, Giessen, Germany), were positioned over the opening above the cortical surfaces on opposite sides of the V1/V2 boundary. Three to six tetrodes were lowered independently into and through the cortex with microdrives (Thomas Recording GmbH, Giessen, Germany) to locations with visually-driven extracellular action potentials. Initial neuronal characterizations were performed to determine tunings for sinusoidal gratings varying in orientation, spatial frequency, temporal frequency, and contrast through on-line analysis of the responses of a few well-isolated single units. The recorded units within either

*MCEN Team, BCAM - Basque Center for Applied Mathematics, 48009 Bilbao, Basque Country, Spain and Department of Mathematics, KTH Royal Institute of Technology, Stockholm, Sweden

†MathNeuro Team, Inria at Université Côte d'Azur, Sophia Antipolis, France

‡Feil Family Brain and Mind Research Institute, Weill Cornell Medical College, New York, NY 10065, USA, (JV)

§Feil Family Brain and Mind Research Institute, Weill Cornell Medical College, New York, NY 10065, USA, (KPP)

¶MCEN Team, BCAM - Basque Center for Applied Mathematics, 48009 Bilbao, Basque Country, Spain and ikerbasque - The Basque Foundation for Science, Bilbao, Basque Country, Spain, (SR)

V1 or V2 generally had overlapping or neighboring receptive fields. The preferred orientation for a cluster of neurons at a recording site was used to choose the orientation for the black-and-white patterns that were used to generate the neural reponses used in this study. Details concerning the procedures for lesion making, perfusion, and histology to identify the recording sites (V1 or V2) and assign laminar location can be found in [1].

### Figures

Eleven supplementary figures (Figs. S1–S11) are included below, on pages 3–13.

### References


[1] Anita M Schmid, Keith P Purpura, and Jonathan D Victor. Responses to orientation discontinuities in V1 and V2: physiological dissociations and functional implications. *Journal of Neuroscience*, 34(10):3559–3578, 2014.

[2] Jonathan D Victor, Daniel J Thengone, Syed M Rizvi, and Mary M Conte. A perceptual space of local image statistics. *Vision research*, 117:117–135, 2015.

[3] Yunguo Yu, Anita M Schmid, and Jonathan D Victor. Visual processing of informative multipoint correlations arises primarily in V2. *eLife*, 4:e06604, 2015.

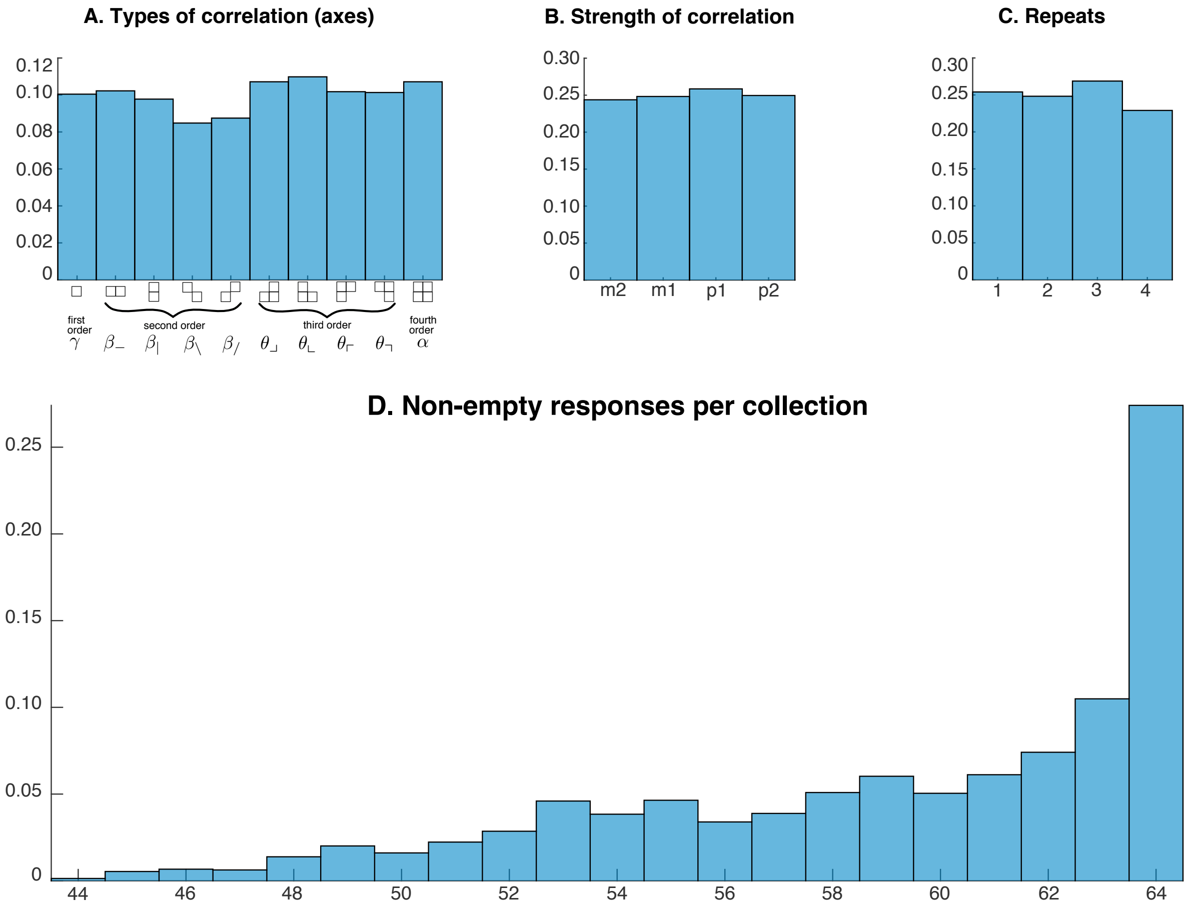


Figure S1: **Analyzed collections of neuronal responses. A,B.** Distributions of the analyzed responses over the 10 axes in the stimulus space that specify the type of spatial correlation and coordinates along those axes that determine the strength of correlation as described in [2]. Compare with Fig. 1 of the main text. **C.** Distribution of the repeats from which the analyzed responses were drawn. **D.** Distribution of the number of non-empty responses (at least one spike) in the $28 \times 80$ collections of 64 responses considered in the analysis.

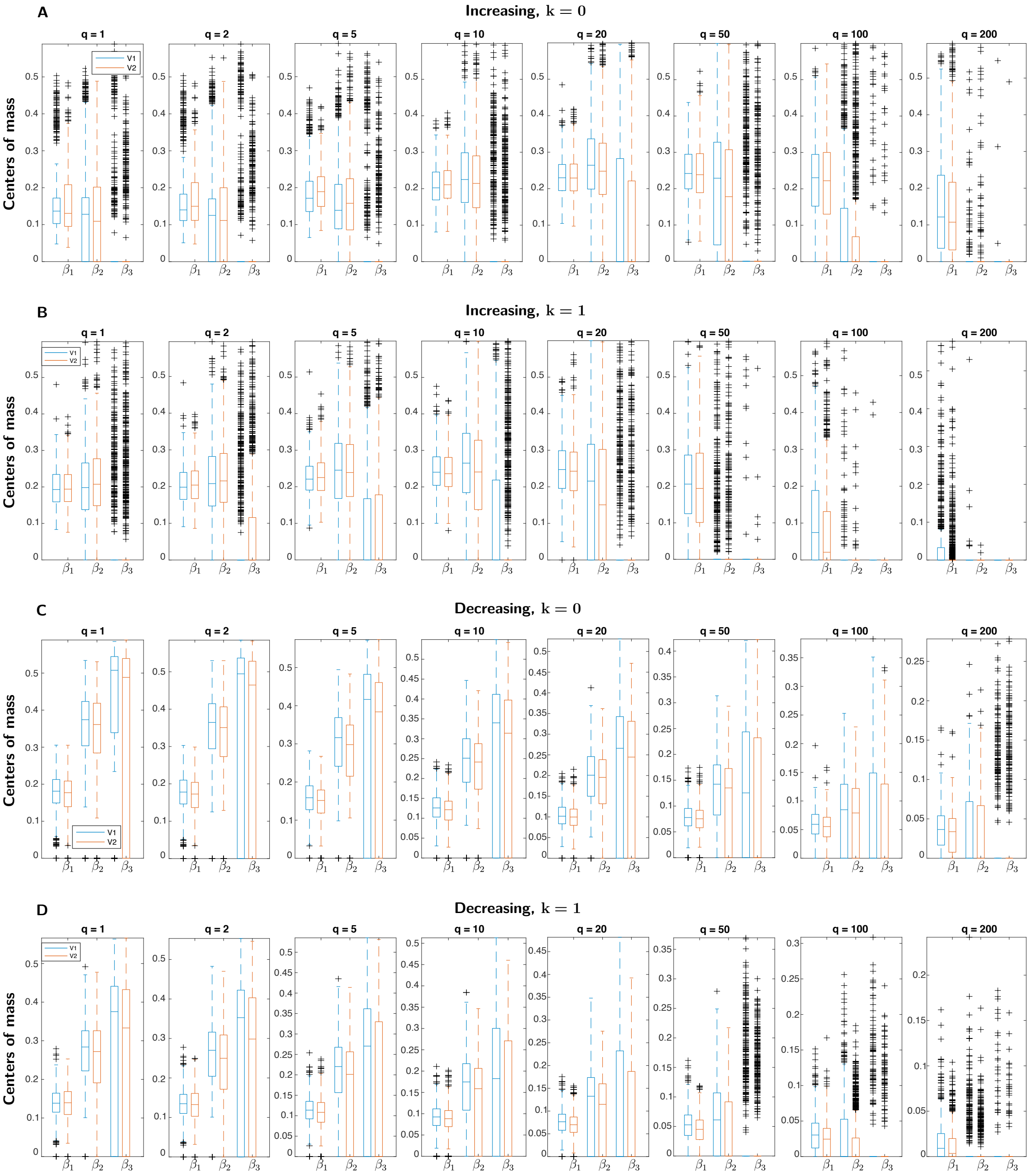


Figure S2: **Comparison of centers of mass of the Betti curves for data recorded in V1 vs. V2.** The figure is similar to Fig. 3 of the main text, but Betti curves are summarized via their centers of mass (see Materials and Methods of the main text) instead of their integrated Betti values. The values of the centers of mass of the Betti curves for $\beta_1$-$\beta_3$ of the experimental data are shown divided into two groups, according to whether the data were recorded in area V1 (blue) or V2 (red) of the visual cortex. To generate the distributions of centers of mass displayed in the figure, individual (non-averaged over the dataset) Betti curves of each collection of responses are considered. The four panels are for increasing (**A**,**B**) and decreasing (**C**,**D**) filtrations, and for $k = 0$ (**A**,**C**) and $k = 1$ (**B**,**D**). Each panel shows the distribution of centers of mass for all values of the timescale parameter $q$ of the Victor-Purpura distance.

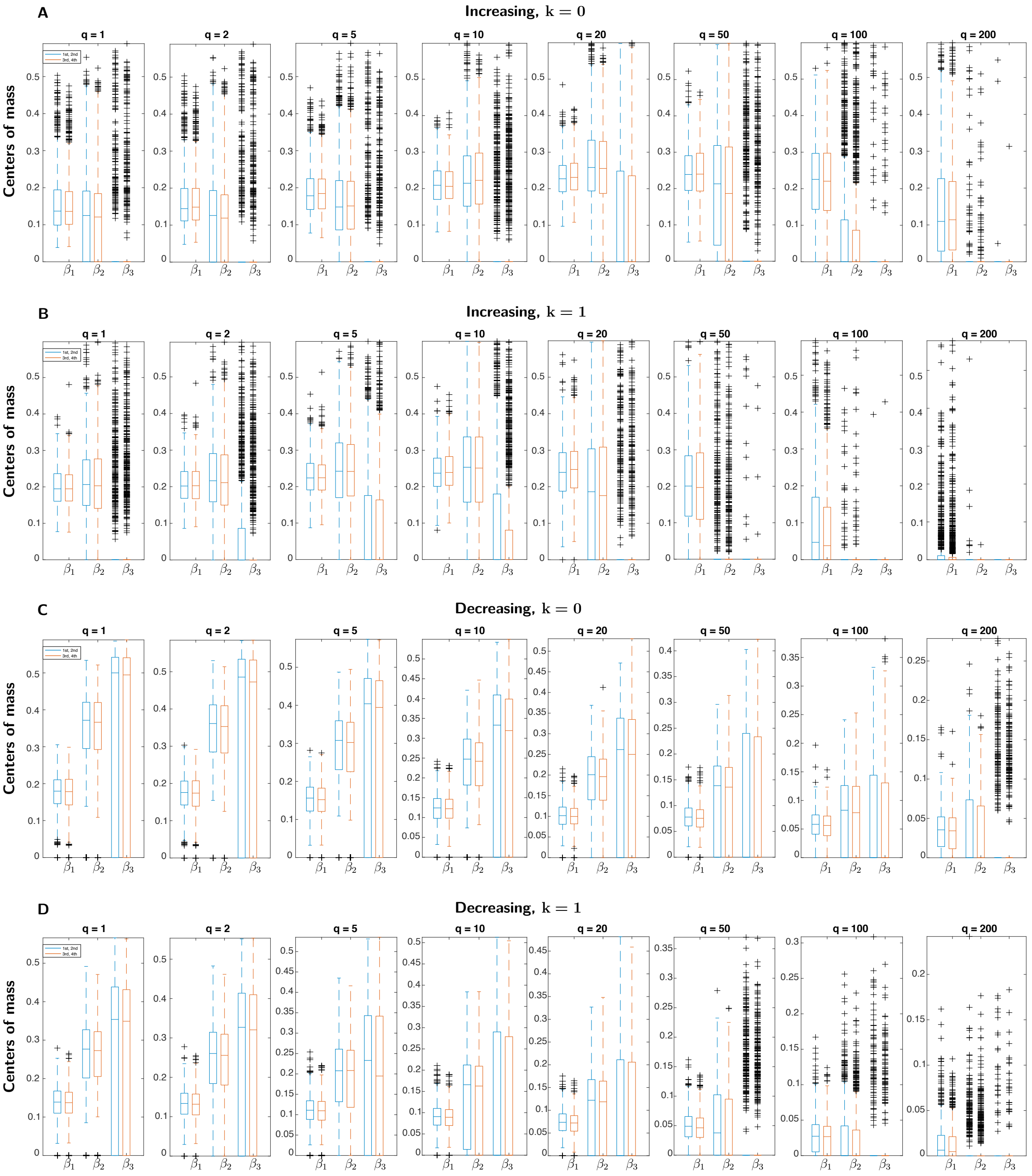


Figure S3: **Comparison of centers of mass of the Betti curves for responses elicited by low- and high-order spatial correlations.** The figure is similar to Fig. 4 of the main text, but Betti curves are summarized via their centers of mass instead of their integrated Betti values. The values of the centers of mass of the Betti curves for $\beta_1$-$\beta_3$ of the experimental data are shown divided into two groups, according to whether neuronal responses are driven by visual stimuli with first- and second-order correlations (axes $\gamma$, $\beta_{-}$, $\beta_{|}$, $\beta_{\backslash}$, $\beta_{/}$ of the stimulus space described in [2] and summarized in Fig. 1 of the main text) or third- and fourth-order correlations (axes $\theta_{\lrcorner}$, $\theta_{\llcorner}$, $\theta_{\ulcorner}$, $\theta_{\urcorner}$ and $\alpha$). The distributions of centers of mass, determined as in Fig. S2, are respectively shown in blue and red. To generate the distribution of centers of mass, individual (non-averaged over the dataset) Betti curves of each collection of responses are considered. The four panels are for increasing (**A**,**B**) and decreasing (**C**,**D**) filtrations, and for $k = 0$ (**A**,**C**) and $k = 1$ (**B**,**D**). Each panel shows the distribution of centers of mass for all values of the timescale parameter $q$ of the Victor-Purpura distance.

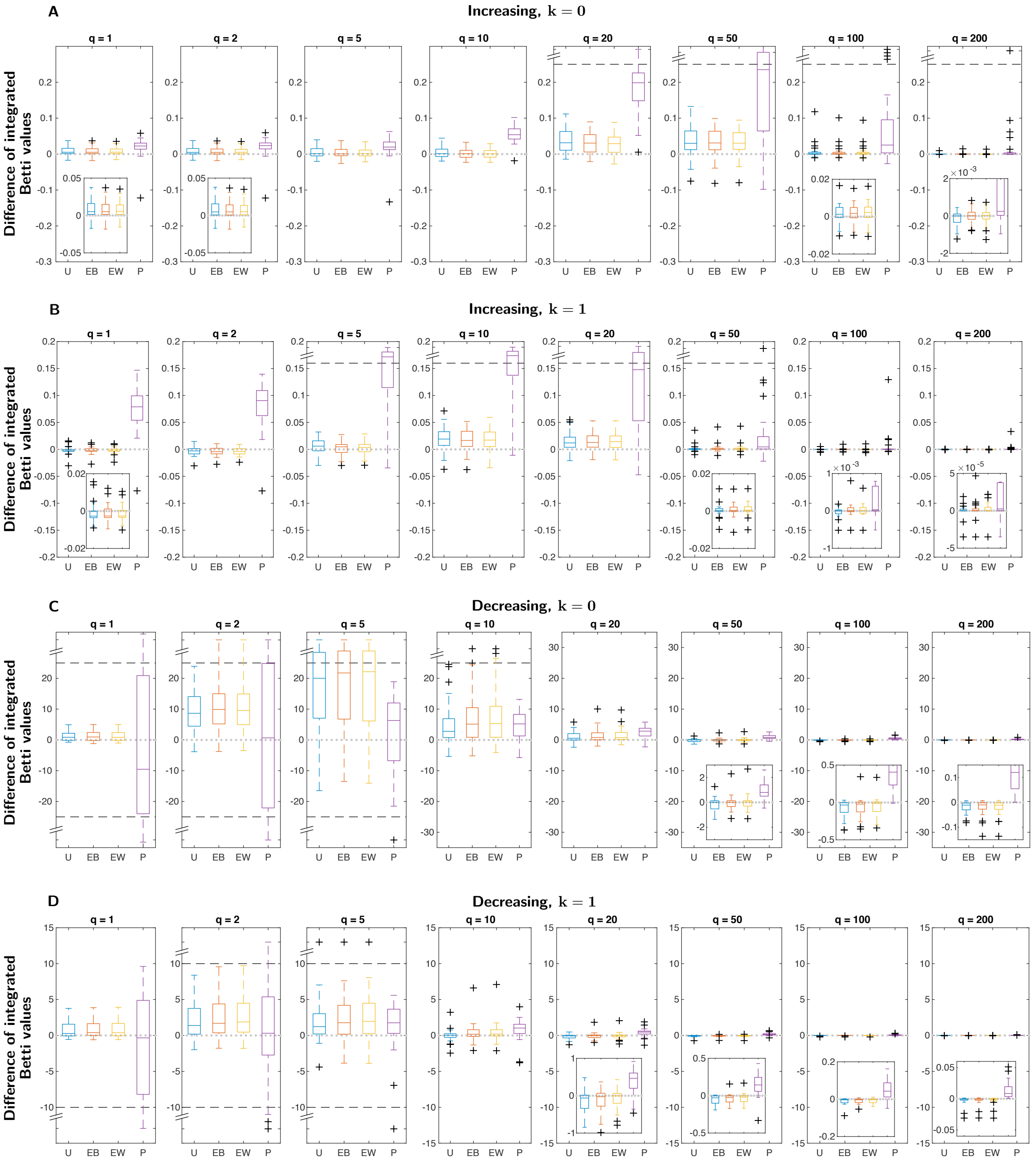


Figure S4: **Distribution of the integrated Betti values for $\beta_2$.** The figure is parallel to Fig. 6 of the main text, but for $\beta_2$ rather than $\beta_1$. It shows the distribution across the 28 datasets of the difference between the integrated Betti values for $\beta_2$ of each surrogate, averaged over the 20 computations, and the integrated Betti value of the experimental data. Each plot shows the four surrogates: uniform resampling of spike times (blue, "U"), exchange of spike times between collections (red, "EB"), exchange of spike times within collections (yellow, "EW"), Poisson generated spike data (purple, "P"). Insets zoom in on some boxplots at a smaller scale. When present, a dashed line bounds an area at the extreme of a plot beyond which data are shown on a compressed ordinate. The four panels are for increasing (**A**,**B**) and decreasing (**C**,**D**) filtrations, and for $k = 0$ (**A**,**C**) and $k = 1$ (**B**,**D**). For example, in **A**, the means of the integrated Betti values for the Poisson surrogates for all timescales, $q$ are greater than the means of the integrated Betti values for the experimental spiking responses from the 28 datasets. Note that the deviation of the behavior of the Poisson surrogates is maximal for the range $q = 5$ sec$^{-1}$ to $q = 50$ sec$^{-1}$.

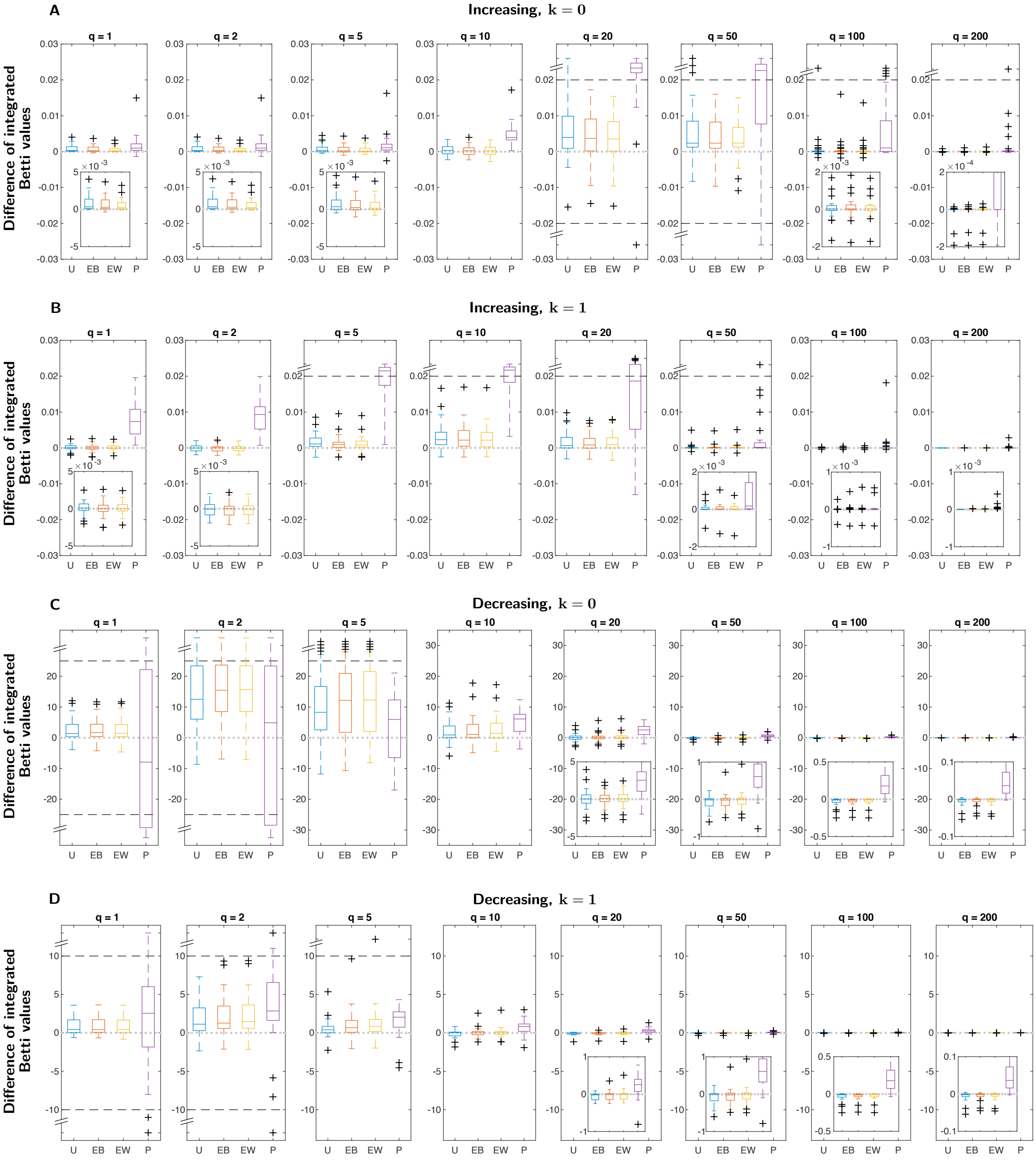


Figure S5: **Distribution of the integrated Betti values for $\boldsymbol{\beta_3}$.** The figure is obtained similarly to Fig. 6 of the main text and Fig. S4, but for $\beta_3$. Plotting conventions as in main text Fig. 6 and Fig. S4.

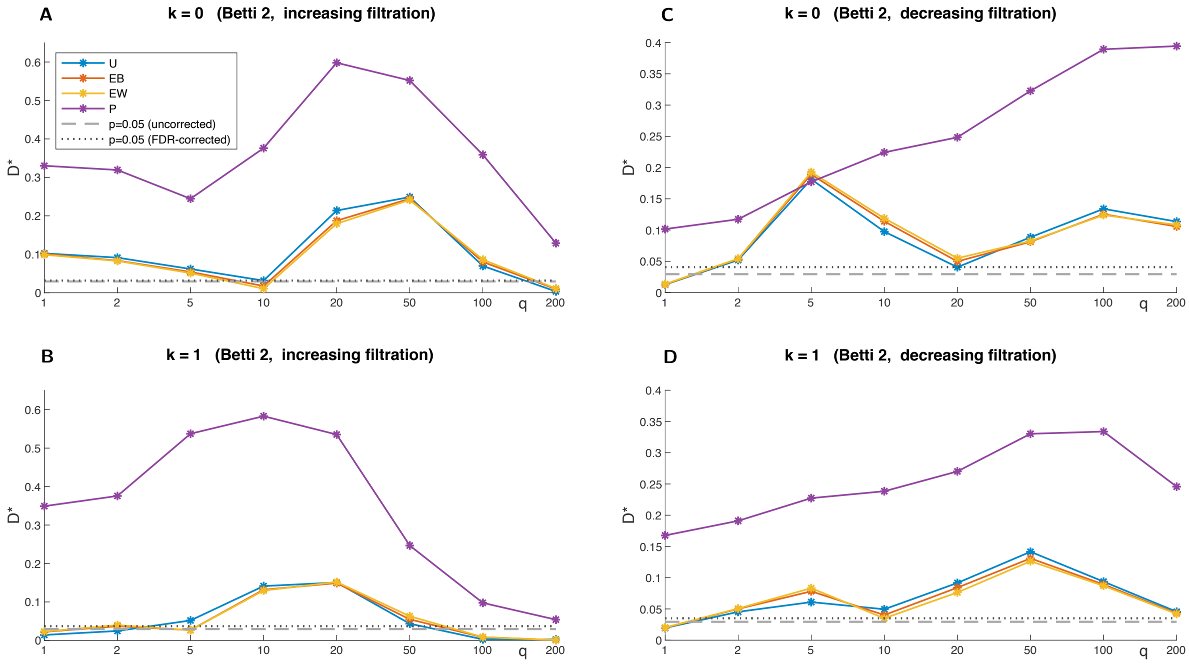


Figure S6: **Two-sample Kolmogorov-Smirnov test for the integrated Betti values for $\beta_2$.** The figure is parallel to Fig. 7 of the main text, but for $\beta_2$ rather than $\beta_1$. It shows the two-sample Kolmogorov-Smirnov test statistic $D^*$ (ordinate) for comparison of the samples of integrated Betti values for $\beta_2$ of the experimental data with each type of surrogate data. The sample for the experimental data consists of the $(28 \times 80)$ integrated Betti values from all 80 collections from all 28 datasets. The samples for each surrogate consist of the $(20 \times 28 \times 80)$ integrated Betti values of the 20 computations. Values above the dashed line, corresponding to $p = 0.05$, indicate rejection of the null hypothesis that the two samples come from the same distribution. The dotted line corresponds to the value $p = 0.05$ corrected for multiple comparisons, using the false discovery rate method. The four panels are for increasing (**A**,**B**) and decreasing (**C**,**D**) filtrations, and for $k = 0$ (**A**,**C**) and $k = 1$ (**B**,**D**).

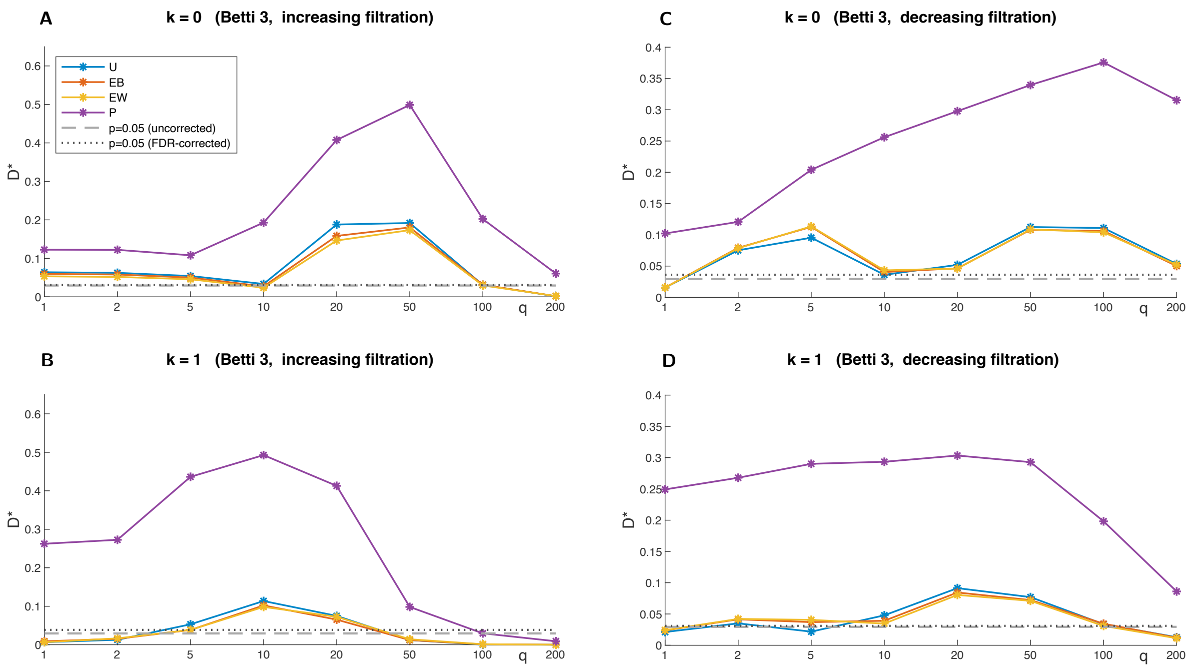


Figure S7: **Two-sample Kolmogorov-Smirnov test for the integrated Betti values for $\beta_3$.** The figure is parallel to Fig. 7 of the main text and Fig. S6, but for $\beta_3$. Plotting conventions as in main text Fig. 7 and Fig. S6.

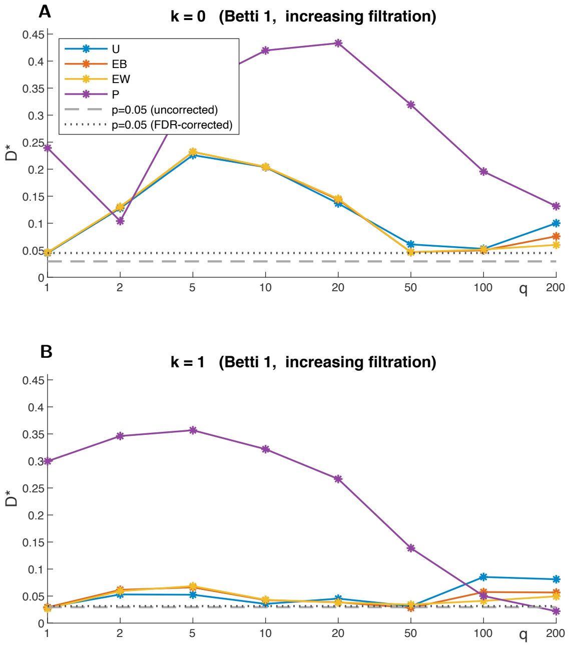


Figure S8: **Two-sample Kolmogorov-Smirnov test for the centers of mass of the Betti curves for $\beta_1$.** Two-sample Kolmogorov-Smirnov test statistic $D^*$ (ordinate) for comparison of the samples of centers of mass of the Betti curves for $\beta_1$ of the experimental data with each type of surrogate data. The sample for the experimental data consists of the $(28 \times 80)$ center of mass values from all 80 collections from all 28 datasets. The samples for each surrogate consist of the $(20 \times 28 \times 80)$ center of mass values of the 20 computations. Values above the dashed line, corresponding to $p = 0.05$, indicate rejection of the null hypothesis that the two samples come from the same distribution. The dotted line corresponds to the value $p = 0.05$ corrected for multiple comparisons, using the false discovery rate method. The four panels are for increasing (**A**,**B**) and decreasing (**C**,**D**) filtrations, and for $k = 0$ (**A**,**C**) and $k = 1$ (**B**,**D**).

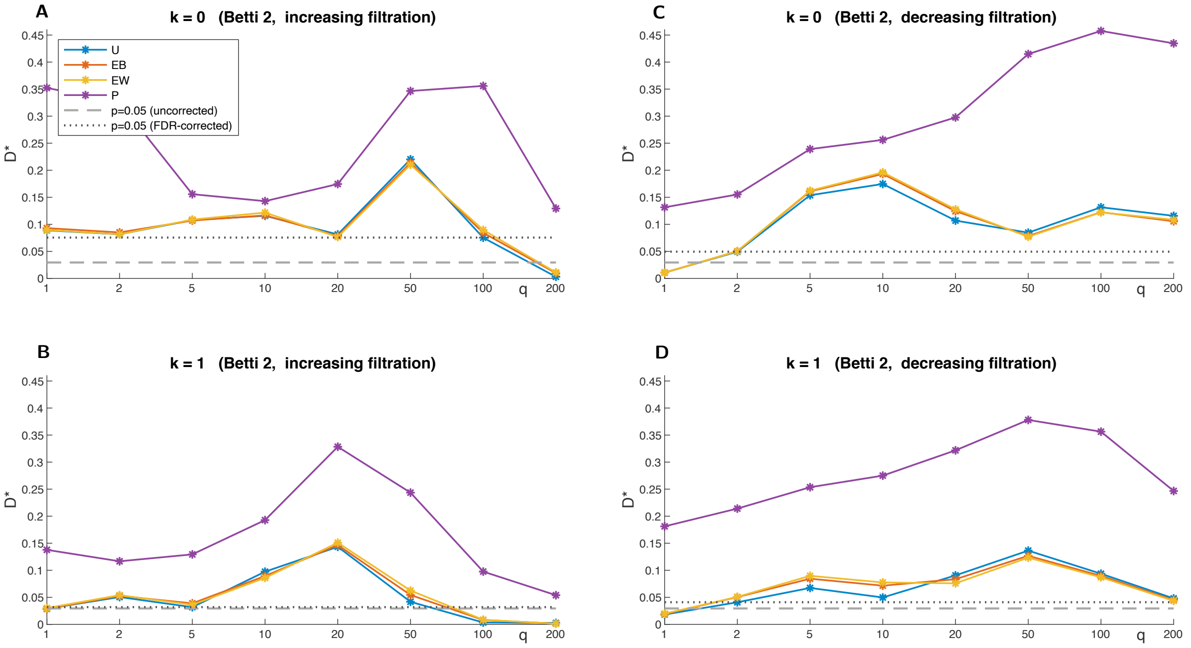


Figure S9: **Two-sample Kolmogorov-Smirnov test for centers of mass of the Betti curves for $\boldsymbol{\beta_2}$.** The figure is parallel to Fig. S8, but for $\beta_2$. Plotting conventions as in Fig. S8.

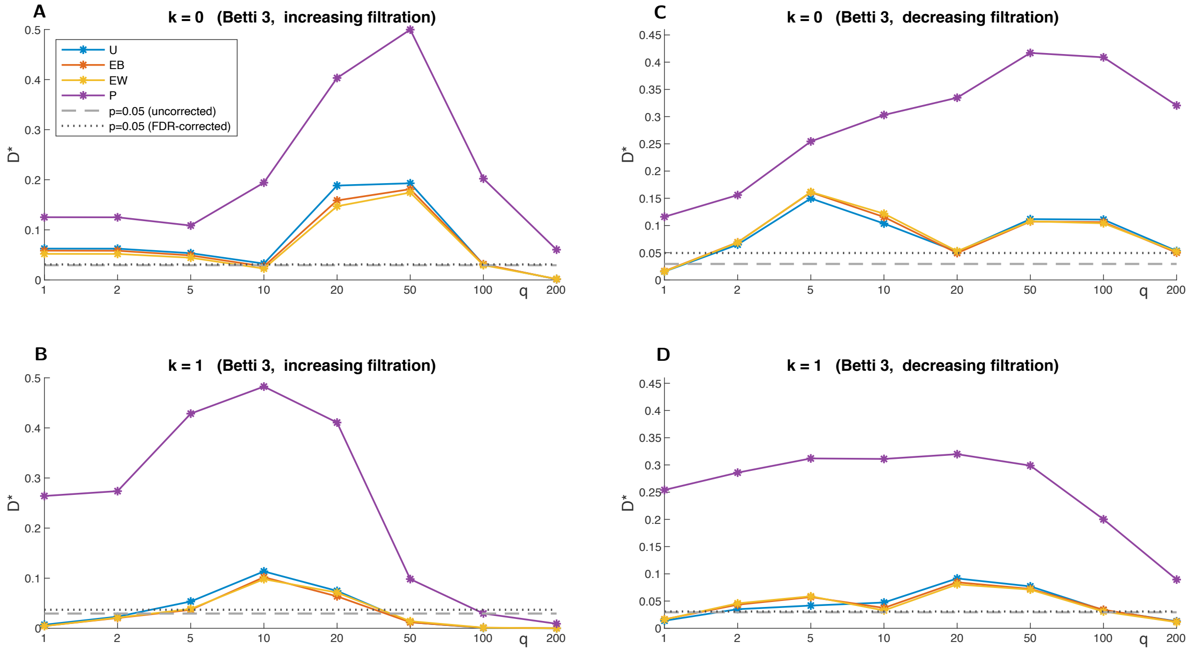


Figure S10: **Two-sample Kolmogorov-Smirnov test for the centers of mass of the Betti curves for $\boldsymbol{\beta_3}$.** The figure is parallel to Figs. S8 and S9, but for $\beta_3$. Plotting conventions as in Fig. S8.

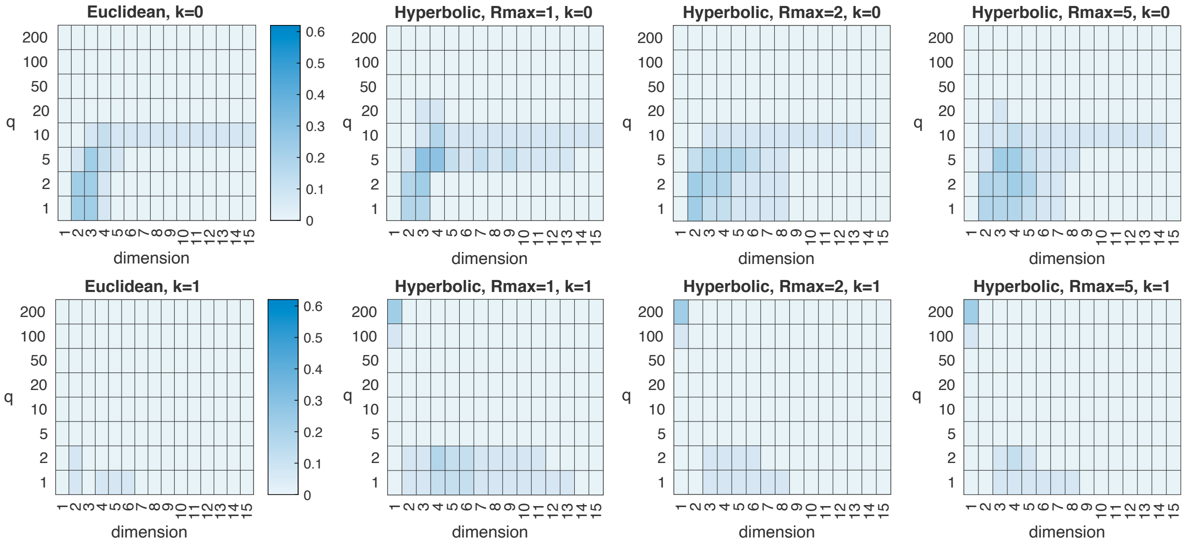


Figure S11: **Compatibility of the experimental Betti curves with geometric models, based on centers of mass of the Betti curves.** The figure is parallel to Fig. 8 of the main text, but compatibility is assessed via the centers of mass of the Betti curves. The heatmaps show the fraction of Betti curves of 18 selected collections from different datasets which are compatible with the Euclidean (column 1) and hyperbolic models ($R_{\max} = 1, 2, 5$) (columns 2-4) of dimension $d = 1, \ldots, 15$ (abscissas), for all values of the parameters $q$ (ordinates) and $k$ (rows) of the Victor-Purpura distance. The notion of compatibility we introduced requires the experimental centers of mass of the Betti curves to be within 3 standard deviations of the mean of the 300 values of the model, for all Betti numbers $\beta_1$-$\beta_3$ and for both increasing and decreasing filtrations. In the range $q = 1$ to 10 sec$^{-1}$, the greatest compatibility occurs for dimensions 2-4. Compatibility is consistently low for $k = 1$ (bottom row), and is generally lower than compatibility assessed via integrated Betti values (Fig. 8 of the main text). As we determined by visual inspection of the Betti curves, the hotspot at dimension 1 and $q = 200$ in the hyperbolic heatmaps for $k = 1$ is an artifact due to the fact that the low dimension constrains the Betti curves of the hyperbolic models to being close to (identically) zero, hence compatible in some cases with the experimental Betti curves at the extreme value $q = 200$.
